# An optimized workflow for analyzing extracellular vesicles as biomarkers in liver diseases

**DOI:** 10.1101/2023.01.30.526180

**Authors:** Martha Paluschinski, Sven Loosen, Claus Kordes, Verena Keitel, Anne Kuebart, Timo Brandenburger, David Schöler, Marianne Wammers, Ulf P Neumann, Tom Luedde, Mirco Castoldi

**Affiliations:** Department of Gastroenterology, Hepatology and Infectious Diseases, Medical Faculty, Heinrich-Heine University, Düsseldorf, Germany; Department of Anesthesiology, Medical Faculty, Heinrich-Heine University, Düsseldorf, Germany; Visceral and Transplant Surgery, University Hospital RWTH Aachen, Aachen, Germany

**Author notes:** To whom address correspondence: Mirco Castoldi; PhD, Department of Gastroenterology, Hepatology and Infectious Diseases, Medical Faculty, Heinrich-Heine University, Düsseldorf, Germany., Moorenstrasse 5, 40225 Düsseldorf, Germany.

**Keywords:** microRNA, extracellular vesicles, liver diseases, biomarker, Nanoparticle-tracking analysis, cytokines

## Abstract

**Background & Aims:** Extracellular vesicles (EVs) play an important role in intercellular communication, serving as vehicles for the exchange of biological materials and being involved in the regulation of physiological processes. EVs and their associated cargoes are considered a promising source of disease-associated biomarkers. The purpose of this study was to establish an easy-to-use, reproducible, and scalable workflow to efficiently analyze EVs in the context of liver disease.

**Methods:** An optimized workflow was established for the pre-analytical processing and isolation of EVs from plasma and serum. Nanoparticle Tracking Analysis (NTA) was used to characterize circulating EVs in the serum of patients with nonalcoholic fatty liver disease (NAFLD), autoimmune liver disease (AIH), and animal models with impaired liver function. EVs were separated from soluble proteins by an optimized, polyethylene glycol (PEG)-based enrichment protocol. Enriched EVs were either labeled and functionally characterized by monitoring cellular uptake or lysed for biomarker identification.

**Results:** Circulating EVs in the serum of patients with NAFLD or AIH and in different animal models have been characterized by NTA. Here we show that both the quantity and size of EVs in the serum of patients/animal models are significantly different from those of healthy individuals. We show that isolated EVs are functional, and their uptake by acceptor cells can be quantified after fluorescence labelling. Enriched EVs were directly used to analyze RNA biomarkers. Several microRNAs, including miR-15b, -16, -21, -122 and -223, were found to be significantly up-regulated in EVs isolated from the sera of patients with NAFLD and AIH. We show that EVs transport cytokines, and that IL-2, IL-6 and IL-8 were significantly up-regulated in EVs enriched from patients with cholangiocarcinoma (CCA) compared to healthy controls.

**Conclusions:** The workflow presented here represents an accessible and easy-to-use approach that enables the analysis and enrichment of EVs from complex biological fluids and their preparation for functional characterization or downstream analysis. In this study, the levels of several miRNAs were found to be significantly increased in EVs isolated from AIH and NAFLD patients compared with healthy controls.

**Highlights:**

- EVs circulating in crude serum reflect the diseased stage of the donors.
- Enrichment of EVs with the approach presented here efficiently separates soluble proteins from EVs, providing optimal material for further characterization.
- Exosomal markers are present in the EVs-enriched fraction.
- Enriched EVs are intact and are functionally taken up by acceptor cells.
- Enriched EVs are suitable, and have been used for, biomarkers identification both at RNA and protein level.

## Introduction

Extracellular vesicles (EVs) are a heterogeneous group of lipid-enclosed nanoparticles secreted by prokaryotic and eukaryotic cells that are mediators of intercellular communication [1]. EVs carry biomolecules including proteins, metabolites, cytokines, mRNAs, and microRNAs (miRNAs), and have the ability to transport these cargoes to recipient cells over short and long distances. The ability of EVs to be transported by the blood and to deliver their payloads to recipient cells makes them excellent candidates for the delivery of agents in targeted therapies and to monitor disease-associated biomarkers [2]. Despite these promising potentials, the use of EVs in the clinical setting is still limited due to the lack of standardized EVs classification, isolation protocols and analysis methods.

The three most prominent subtypes of EVs are the exosomes (Exo, Ø 30 nm to 150 nm, [3]), microvesicles (MVs, Ø 100 nm to 1 µm), and apoptotic bodies (AP, Ø 1 µm to 5 µm), which have been differentiated based upon their biogenesis, release pathways, size, content, and function [4, 5]. While Exo and their molecular composition have been studied intensively and tumor susceptibility gene 101 (TSG101), tetraspanins (e.g., CD63, CD81), and the 70-kDa heat shock protein 70 (HSP70) are all accepted exosomal markers [6], no specific protein markers have been identified to distinguish between the different types of EVs [3]. As an additional confounding factor, it has been shown that the proteomic profiles of EVs from the same source depend on their isolation method [4], which may be partly explained by the lack of standardization of isolation procedures and markers for differentiating EVs subgroups.

The lack of specific markers, overlapping sizes and physical properties of different types of EVs make the isolation of enriched subpopulations of EVs an extremely challenging task, requiring the combination of different methodological approaches and techniques [7, 8], which are often beyond the reach and scope of most research laboratories, especially those working in clinical settings.

The purpose of this study was to establish an easy-to-use, reproducible, and scalable workflow to efficiently analyze EVs in the context of liver disease. Here we show that EVs distribution in sera of patients with nonalcoholic fatty liver disease (NAFLD) or autoimmune hepatitis (AIH) is significantly different from the EVs circulating in the sera of healthy donors. Significant differences were also found in the EVs distribution in an animal model for hyperbilirubinemia (Gunn rats, [9]) as well as in the sera of aging rats. These data indicate that the direct analysis of EVs in serum (or plasma) can potentially provide diagnostic or prognostic information on the health status or age of the donor.

We demonstrated that an optimized mixture of polyethylene glycol (PEG) and sodium chloride (NaCl) is effective in separating lipid-encapsulated nanoparticles from soluble proteins, which remain in the supernatant. Isolated EVs, which are enriched in exosomal markers, can then be used directly for biomarker discovery (e.g., microRNAs and cytokines), or in functional studies, such as monitoring cellular uptake in-vitro. In addition, PEG-enriched EVs can serve as an entry point for further downstream procedures aimed at the enrichment of specific subtype of EVs or in preparation for proteomics analysis of EVs-associated proteins.

## Materials & Methods

### Quantification of EVs by using Nanoparticle Tracking Analyzer and Flow Cytometry

Analysis of EVs was performed with ZetaView multi parameter Particle Tracking Analyzer (ParticleMetrix, Germany). Samples were diluted in PBS to achieve a particle count in the range of 1 – 9 × 10^7^ p./mL (or 250 to 300 particles per visual field, PVF). Using the script control function, five 30-second videos for each sample were recorded, incorporating a sample advance and a 5-second delay between each recording. For analysis of bilirubin (Br) and hemoglobin (Hb), samples were diluted in PBS according to the same dilution factor as sera from control animals (i.e., 1:7000). Bilirubin (Br) stock was prepared in chloroform to a final concentration of 10 mg/mL. For estimating the amount of hemolysis, it was considered that red blood cells (RBCs) circulating in a healthy human adult contain 150 – 200 mg hemoglobin per mL of blood (or 15 – 20 g/dL). Human hemoglobin (Sigma) was dissolved in PBS to 10 mg/mL or 1 mg/mL. For achieving 100 mg/mL, Hb stock was diluted 1:700 (instead of 1:7000) in PBS before NTA measurement. Next, to mimic hemolysis, 0.5% (1 mg/mL), 5% (10 mg/mL) or 50% (100 mg/mL) Hb dilutions were prepared in PBS.

In order to visualize EVs by Flow Cytometer (FC), sera samples and conditioned tissue culture medium from rat hepatic stellate cells (HSCs) or primary rat hepatocytes (PCs) were stained with phycoerythrin (PE)-labeled Cell Mask (Thermo Fisher Scientific) or stained with Syto RNASelect (Thermo Fisher Scientific), measured in FITC channel. Using a BD FACSAria III Cell Sorter (BD Biosciences) equipped with four air-cooled lasers at 375-, 488-, 561-, and 633-nm wavelengths, a 1.0 neutral density filter in front of the forward scatter detector was used to decrease the forward scatter (FSC) signal. To avoid exclusion of the smallest events an absolute minimum threshold of 200 was set at the side scatter (SSC)-A parameter (instead of FSC-A). To locate the size of 200 nm EVs, a gate was established using size-calibrated and FITC-labeled beads with sizes of 200 nm (TetraSpeck, Thermo Fisher Scientific, T7280). Samples were evaluated with Flowjo v 10 (FlowjoLLC, BD).

### Isolation of extracellular vesicles with PEG and TEI, and labeling

Before precipitation, serum samples were precleared by centrifugation at 10,000 relative centrifugal force (rcf) for 30 min at 4°C. Supernatants were transferred to a new tube and sera were either left untreated (CTRL) or diluted with either the recommended amount of TEI [Total Exosome Isolation Reagent from serum (Invitrogen, 4478360)] or different amounts of PEG precipitation solution (PPS). In the initial testing, different amounts of stock solution (24% PEG-8000, 1.5 M NaCl) were added to the sera to obtain final amounts of PEG 4%/ 250 mM NaCl (1:6 dilution), PEG 8%/ 500 mM NaCl (1:3 dilution) and PEG 12%/ 750 mM NaCl (1:2 dilution).

The samples were incubated 30 min at 4°C and then centrifuged (10,000 rcf for 30 min at 4°C). The vesicle-free supernatants were transferred to fresh tubes for NTA or Western blot analysis, while the vesicle-enriched pellets were rinsed twice with 1 mL of cold PBS and finally dissolved in PBS for downstream procedures.

For labeling of EVs, 1 µL of either Cell Mask orange plasma membrane stain (Thermo Fisher Scientific, C10045) or Syto RNASelect (Thermo Fisher Scientific, S32703) was added to 500 µL of serum and incubated 15 minutes at room temperature (RT) before treatment with Proteinase K (Carl Roth, 7528.1) or Triton X-100 or the addition of precipitation reagents. For digestion/lysis EVs-enriched pellets were dissolved in PBS and either incubated with 200 µg/mL of proteinase K for 1 hour at 48°C or with 0.3% Triton X-100 for 15 minutes at RT. Labeled/digested EVs were precipitated with PEG as described above.

### Western blot analyses and antibodies

Protein concentrations were measured using Qubit Protein Assay Kit (Thermo Fisher Scientific, Q33211) according to the manufacturer’s instructions. Western Blot analyses were carried out using 25 µg of total proteins. Serum proteins were first precipitated by addition of 10% (w/v) trichloroacetic acid (TCA; Sigma, 49010) and incubated at -20°C for up to 30 min. Denatured proteins were pelleted by centrifugal force (10 min, 4°C at 10,000 rcf) and washed shortly with cold acetone. Following centrifugation (10 min, 4°C at 10,000 rcf), acetone was removed and the protein pellet was shortly allowed to dry. The proteins were then redissolved in loading buffer (250 mM Tris pH 6.8, 10% (v/v) glycerol, 2% SDS, 0.01% (w/v) bromophenol blue, 50 mM DTT). Protein lysates were loaded together with 10 µL of Protein Ladder (BioRad, 161-0373) on 10% SDS polyacrylamide gels and transferred to nitrocellulose membranes using semidry blotting systems according to standard protocols. Membranes were blocked with 5% milk powder (Carl Roth, T145.3) in Tris-buffered saline with Tween20 (TBST) for 1 hour at RT, followed by incubation with a horseradish peroxidase (HRP)-coupled antibody against rat albumin (Bethyl Laboratories, A110-134P) for 2 h at room temperature. Chemiluminescence was detected with ECL Western Blotting Substrate (Promega, W1001) using the ChemiDoc MP Imaging System (BioRad). Signal intensities of Western blot protein bands were analyzed using ImageJ 1.51 software (National Institute of Health, USA). Antibodies used in protein analysis by Western blot: anti-Albumin antibody (1:2,000 dilution, Abcam, ab207327), anti-TSG101 antibody (1:1,000 dilution, Abcam, ab133586), anti-CD63 antibody (1:200 dilution, Biorad, MCA4754GA), and anti-Hsp70 antibody (1:500 dilution, Cell Signaling, 4872T).

### RNA isolation, visualization and miRNA analysis by miQPCR

EVs were enriched by PEG precipitation starting from 100 µL serum and dissolved in 100 µL of PBS. RNA was isolated by using QIAzol Lysis Reagent (Qiagen, 79306) and the miRNeasy Mini Kit (Qiagen, 217004). cDNA synthesis was carried out by using equal volumes of elute material instead of ng of RNA (as described in [10]). Electropherogram was generated with the Agilent 2100 Bioanalyzer. For this, EVs were enriched from 500 µL of sera, and RNAs were isolated from both EVs-enriched pellets and EVs-depleted sera with the miRNeasy Mini Kit (Qiagen). RNAs were eluted from the miRNeasy column in 100 µL of RNA-free water, and 1 µL of the resulting material was loaded and run on a PicoChip according to the supplied protocol.

miRNA expression profiling by qPCR was performed by using the miQPCR method [10]. qPCR reactions were carried out on a StepOnePlus Real-Time PCR system (Applied Biosystems), whereas amplicons were detected using SYBR Green I (Promega, A6002). Data analysis was carried out by using qBase [11], GeNorm was used to identify appropriate reference genes. Gene expression was normalized to Let-7a, Let-7c and Let-7e by using the ΔΔCt method [12]. Sequences of the miQPCR primers designed to amplify miRNAs included in this study are listed in Supplementary Table 4.

### Preparation of primary cells from rat liver and liver samples from Gunn and aging rats

Primary hepatocytes were isolated from the livers of 2 months-old male Wistar rats (150 – 200 g) essentially as described in [10]. In brief, hepatocytes were isolated after serial perfusion of rat liver by Hanks ‘ balanced salt solution (HBSS, Sigma, H6648) and collagenase CLS type II solution (50 mg/150 mL, Biochrom, C2-22). Following digestion and centrifugation, pelleted hepatocytes were resuspended in culture medium (Williams ‘ E medium, Millipore, F1115) supplemented with 10% (v/v) fetal calf serum (FCS superior, Millipore, S0615), 2 mM *L*-glutamine (Gibco, 25030), 0.6 µg/mL insulin (Sigma, I05116), 100 nM dexamethasone in DMSO (Sigma, D8893) and 1% (v/v) penicillin-streptomycin-amphotericin B solution (Gibco, 15240-062). Hepatocytes were seeded on polystyrene tissue culture dishes of appropriate size pre-coated by collagen type I (Sigma, C3867, 6 – 10 mg/cm^2^). The homozygous mutant Wistar rat strain (Gunn-UGT1A1; [13]) lacking the uridine diphosphate glucuronosyltransferase-1A1 (UGT1A1) enzyme activity and exhibiting elevated bilirubin serum levels were used in this study. Total bilirubin concentration of the Gunn rat serum was determined by the central hospital laboratory of Heinrich Heine University. Gunn rats were obtained from the Rat Resource & Research Centre (RRRC, Columbia, MO, USA) and maintained at the animal facility of the Heinrich Heine University (Düsseldorf, Germany). Normal Wistar rats were obtained from Janvier Labs (France) for liver aging studies. Young (2-months-old) and aged rats (22-months-old) from the same breeding colony with matching housing conditions were used to obtain blood serum samples after cannulation of the portal vein.

### Visualization and quantification of EVs uptake

Primary hepatocytes isolated from rat livers were plated in 24-well plates coated with collagen type I in William ‘s E (WE) medium supplemented with 10% FCS and were allowed to attach and recover for 24 hours. Thereafter, cells were incubated in serum-free WE medium containing ∼2.0 × 10^11^ of fluorescently labeled EVs with/without Proteinase K (PTK) digestion or negative control (i.e., mock labeling) and cultured for 24 hours at 37°C and 5% CO_2_ in humidified atmosphere. Then, the culture medium was exchanged, and the cells were incubated for further 24 hours. Images of live cells were taken with an Olympus IX50 fluorescence microscope equipped with a DP71 digital camera (Olympus) using the excitation filter set 470/22 nm and the emission filter set 510/42 nm.

For the quantification of EVs uptake, background fluorescence was assessed by measuring fluorescence before the addition of labeled EVs (Time 0). Following the addition of EVs, a measurement was taken every 90 minutes for 5 times. Fluorimetric quantification was carried out by microplate fluorimeter (Fluoroskan Ascent FL) after excitation at 485 nm. The emitted fluorescence light was measured at 538 nm. Data were analyzed by using Microsoft Excel and GraphPad Prism.

### Patient ‘s material and cytokine quantification by Luminex

NAFLD and AIH samples included in this study were from patients visiting the hepatitis outpatient clinic of the University Hospital of Düsseldorf (i.e., not hospitalized). The hepatitis outpatient clinic is a trans-regional center for patients with chronic viral diseases of the liver. Only sera of patients negative to viral hepatitis were included in this study. Healthy volunteers from our department donated the blood for the control group. Analysis of liver parameters was carried out by the central hospital laboratory of the Heinrich-Heine University.

The CCA cohort consists of n=16 patients with intrahepatic cholangiocarcinoma (CCA) who were admitted to the Department of Visceral and Transplantation Surgery at University Hospital RWTH Aachen for tumor resection and were recruited between 2011 and 2017 (see Supplementary Table 5 for patient characteristics). Whole blood samples were taken before surgery, centrifuged for 10 min at 2,000 g, and serum samples were then stored at -80°C until use. Billiary tract cancer (BTC) was confirmed histologically in the resected tumor sample. For the preparation of serum from healthy controls, whole blood samples were taken at RT, and collected into Sarstedt collection tubes (serum: S-Monovette 7,5 mL Z / plasma: S-Monovette 2.7 mL K2 EDTA, both Sarstedt, Nümbrecht, Germany) and were centrifuged for 10 min at 2,000 rcf. The supernatant (serum or plasma) was carefully transferred into a fresh tube and stored at -80°C until use.

EVs isolation for Luminex quantification of cytokines in sera from the CCA cohort was carried out by using PEG 4%/ 250 mM NaCl (1:6 dilution) as described above. To ensure a complete removal of serum proteins, the procedure was repeated three times. Following the third precipitation, EVs were diluted in 25 µL of lysis buffer (0.3% Triton X-100 in PBS) and incubated for 15 min at RT. Samples were analyzed by multiplex immunoassay according to the manufacturer ‘s instruction using a Bio-Plex 200 system and Bio-Plex Manager 6.0 software on a Bio-Plex Pro Human Chemokine Panel (Bio Rad, 171AK99MR2) including the cytokines: IL-2, IL-4, IL-6, IL-8, IL-10, GM-CSF, INFγ and TNFα. A serum sample volume of 50 µL, or an amount of enriched EVs equivalent to 50 µL of serum, were used.

### Statistical analysis

Statistical analyses were carried out by using GraphPad Prism. To evaluate whether samples were normally distributed, D ‘Agostino and Pearson normality test was carried out. When samples distribution passed the normality test, then parametric tests were carried out (i.e., One Way analysis of variance/ANOVA for three, or more, samples and two tailed T-test for two samples). When the samples did not pass the normality test, then non-parametric tests were applied (i.e., Kruskal-Wallis test for three or more samples and Mann-Whitney test for two samples). Data were considered significant at a P value ≤ 0.05.

### Ethical Statement

The ethics committee of the University of Düsseldorf and the relevant federal state authority for animal protection (Landesamt für Natur, Umwelt und Verbraucherschutz Nordrhein-Westfalen, Recklinghausen, Germany) specifically approved the isolation protocol for rat hepatocytes (reference no. 82-02.04.2015.A287 and 81-02.04.2018.A126). All patients were consented at the University hospital of Düsseldorf and the ethics committee of the University of Düsseldorf approved the collection of the samples. The study was performed according to the guidelines of the Declaration of Helsinki, approved by the local ethics committee and informed written consent was obtained from each patient (reference no. 5350 and 3115).

The study protocol for the CCA cohort was approved by the ethics committee of the University Hospital RWTH Aachen, Germany (EK 206/09) and conducted in accordance with the ethical standards laid down in the Declaration of Helsinki. Written informed consent was obtained from all patients.

## Results

### Characterization of Extracellular Vesicles in crude sera

Emerging evidence suggest that the quantity and composition of secreted vesicles change in different pathological conditions [14, 15]. Hence, the accurate characterization of EVs is essential to obtain further knowledge on their biological relevance. Several techniques are available to characterize EVs [7, 16], including flow cytometry (FC), nanoparticle tracking analysis (NTA), transmission electron microscopy (TEM) and dynamic light scattering (DLS). In order to study the impact of liver diseases on the distribution of secreted EVs, NTA was used in this study to monitor and compare EVs characteristics. NTA is a technique based on light-scattering, which enables the rapid sizing and enumeration of nanosized particles in solution [17]. Since its introduction, NTA has become a standard procedure to reproducibility measure EVs size and concentration in their native conditions [18, 19]. Importantly, precision and reproducibility of NTA measurements was carried out by using standardized procedure (Supplementary Figure 3, Supplementary Figure 4, and Supplementary Table 2).

In order to evaluate the performance of the NTA in different experimental models, EVs size and concentration was measured in the sera of human patients with NAFLD, AIH and healthy donors, in sera isolated from young and aged rats and in the sera obtained from Gunn rats with homozygote *Ugt1a* mutation and control rats (Figure 1, Representative NTA images are shown in Supplementary Figure 1).

**Figure 1.**
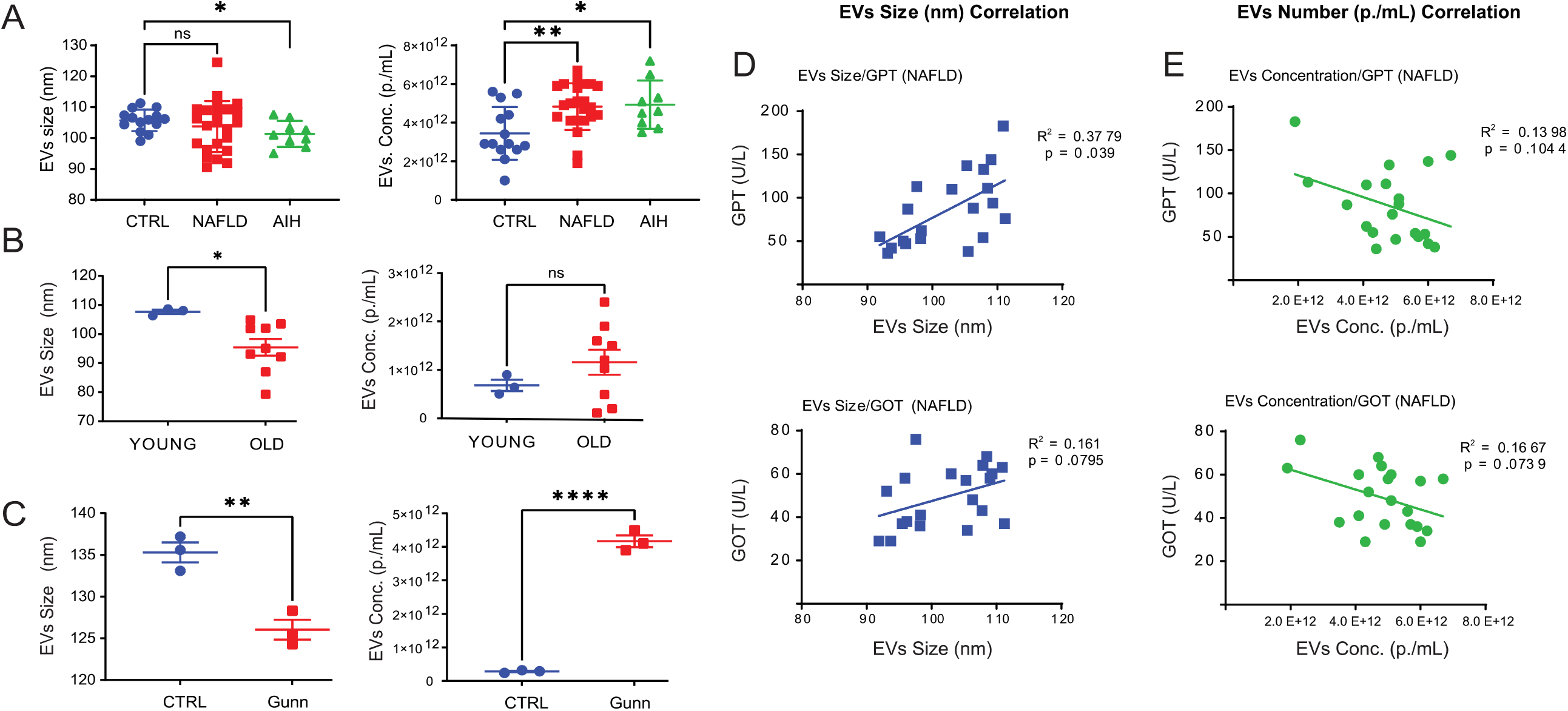
NTA quantification of EVs in sera of liver-diseased patients. Sera from patients with either NAFLD (n = 24), or AIH (n = 9) or healthy donors (n = 14) were analyzed by NTA. (**A**) Average particle size as measured by NTA showed a reduced EV size in AIH patients compared to healthy individuals, while quantification of EVs in patients ‘ sera indicated an increased number of EVs in NAFLD and AIH patients compared to control, respectively. (**B– C**) Analysis of EVs size distribution and number in the sera of (**B**) younger (2-months-old) and older (22-months-old) rats (n=3 and n=9, respectively) and in the sera of (**C**) Gunn and control rats (n=3). Correlations of serum GOT and GPT, with particle average sizes (**D**) and EV number (**E**) in NAFLD patients. Statistical analyses were carried out in GraphPad using one-way ANOVA or two tailed unpaired t-tests for comparison of two groups. p ≤ 0.05, *; p ≤ 0.01, **; p ≤ 0.001, ***; p ≤ 0.0001, ****.

Here we show that vesicles circulating in patients ‘ sera were significantly smaller (CTRL vs. AIH, p = 0.0128) and more abundant (CTRL vs. NAFLD, p =0.0027; CTRL vs. AIH, p =0.0155) compared to EVs present in the sera of the healthy donor group (Figure 1A and Supplementary Table 2). Notably, a similar reduction of EVs size (young vs. old rats size, p =0.0378) and a trend toward increased in EVs number was observed also in sera prepared from aging rats. Moreover, a similar trend was observed in animal model for hyperbilirubinemia [9] (Supplementary Table 1), which also displayed smaller (CTRL vs. Gunn rats, size, p =0.0053) and more abundant EVs (CTRL vs. Gunn rats, number, p <0.0001).

To determine whether the distribution (e.g., size and number) of EVs circulating in patients with NAFLD and AIH correlates with liver injury, EVs-parameters were compared with the available baseline of liver enzymes measured in the patients ‘ sera (Table 1), but no significant correlation was found for the AIH patients (data not shown). Regarding the NAFLD patients, a significant positive correlation was found between the levels of glutamyl transpeptidase (GTP) and EVs size (R^2^ = 0.3779, p = 0.039, Figure 1D), and a weak positive correlation was found between the levels of glutamic-oxaloacetic transaminase (GOT) and EVs size (R^2^ = 0.161, p = 0.0795, Figure 1E). Interestingly, a weak negative correlation was observed between GTP (R^2^ = 0.1398, p = 0.1044, Figure 1F) and GOT (R^2^ = 0.1667, p = 0.00739, Figure 1G) and Evs number.

**Table 1.** Baseline biochemical parameters in NAFLD and AIH patient groups

**Fig.6**
| Table 1 | NAFLD (n = 24) |  | AIH (n = 9) |  | Statistical Analysis |
| --- | --- | --- | --- | --- | --- |
|  | Range | Mean±SD | Range | Mean±SD |  |
| GOT (U/L) | 23 - 76 | 44.3±16.3 | 27 - 337 | 98.2±98.9 | n.s. |
| GPT (U/L) | 12 - 144 | 65.7±41.9 | 29 - 451 | 120.1±130.6 | n.s. |
| GGT (U/L) | 8 - 657 | 132.5±156.3 | 14 - 589 | 201.6±173.4 | n.s. |
| AP (U/L) | 37 - 195 | 75.5±34.6 | 60 - 158 | 108.1±32.0 | p = 0.0037 |
| Albumin | 4.0 - 5.1 | 4.56±0.31 | 4.0 - 4.6 | 4.29±0.26 | p = 0.0153 |
| Bilirubin (mg/dL) | 0.17 - 2.51 | 0.55±0.48 | 0.27 - 0.58 | 0.44±0.10 | n.s. |

### Assessing the effect of bilirubin, hemoglobin and proteins aggregates on NTA measurements

Due to the principles through which NTA achieves the detection and quantification of particles in suspension, NTA-detected signals in complex biological fluids could be generated either by cell-derived nano-vesicles (i.e. EVs and lipoproteins) or by other structures including aggregates of serum proteins. NTA may also detect the presence of autofluorescent proteins such as bilirubin (Br) or hemoglobin (Hb).

EVs are readily lysed by the addition of detergent such as Triton X-100 [20]. Hence, to assess whether protein aggregates are present and detectable in serum, sera were incubated with either Proteinase K (PTK), or Triton X-100 (TX100), a combination of both (PTK/TX100), or left untreated. Samples were incubated either at room temperature (CTRL and TX100) or at 48°C (CTRL-48°C, PTK and TX100/PTK). The effect of PTK and TX100 treatments on proteins was assessed by Western blot (Figure 2A). Here we show that PTK-mediated digestion of proteins did not significantly affect the number nor the size of EVs as measured by NTA (Figure 2B). However, TX100 alone or in combination with PTK administration did result in the partial (TX100) or complete (TX100/PTK) removal of signal detected by NTA compared to both, control sera and PTK digested sera.

**Figure 2.**
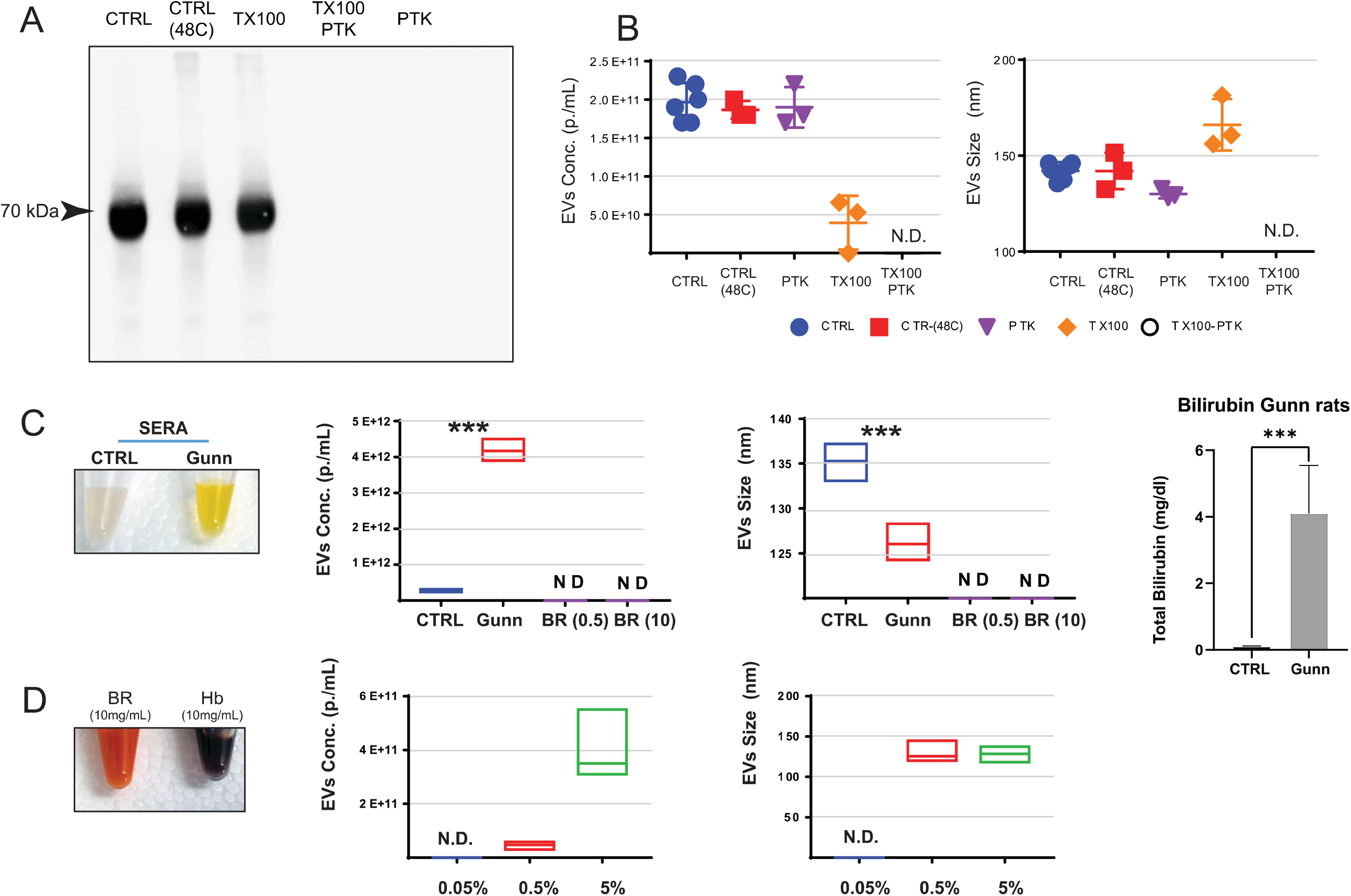
(**A**) Western Blot for albumin in rat sera treated with either 500 µg/mL Proteinase K (PTK), 1% Triton X-100 (TX100), or the combination of the two (PTK/TX100) and incubated at 48°C overnight. Control samples were left untreated (CTRL) or incubated at 48°C overnight [CTRL(48C)]. The figure shows protein bands with 70 kDa size (integration time 50 s). (**B**) NTA analysis of the PTK, TX100, PTK/TX100 treated or control sera (n = 6). (**C**) EVs circulating in sera from Gunn and control (CTRL) rats were analyzed by NTA. EVs concentration (left panel) and size (right panel) in the sera of CTRL and Gunn rats, showing a significant increase of EVs concentration and reduction in EVs size in Gunn compared to CTRL rats (n=3). To evaluate the potential effects of Bilirubin (Br), NTA measurement in crude rat serum (or plasma), Br was diluted at 0.5 mg/mL [BR(0.5),] and 10 mg/mL [BR(10)] in PBS (see material and method section) and measured by NTA. No signal was detected by NTA in Br solutions. (**D**) To evaluate the potential effects of hemolysis of NTA, hemoglobin (Hb) was diluted in PBS to simulate 0.5% and 5% hemolysis (see material and method), and measured by NTA. Data are shown as average ± standard deviation (n = 5), statistical analyses were carried out in GraphPad using one-way. p ≤ 0.001, ***.

In healthy individuals Br, a product of heme catabolism, is present at concentrations below 2 mg/dL. However, in individuals affected by biliary obstruction or by Crigler-Najjar syndromes, Br can rise up to 20 mg/dL (or 340 μmol/L, [21]). One of the animal models for the Crigler-Najjar syndrome is the Gunn rat, which closely parallels the genetic lesion observed in humans [9]. Analysis of EVs in Gunn rats ‘ sera identified significant differences in both the number (p < 0.0001; Gunn; 4.17×10^12^ ± 1.76×10^11^ vs. CTRL; 2.83×10^11^ ± 2.33×10^10^) and the size (p < 0.005; Gunn; 126 ± 1.18 nm vs. CTRL; 135.3 ± 1.19 nm) of EVs (Figure 1C). To assess whether the observed differences were disease-driven or due to higher-than-normal levels of Br, solutions containing different amount of Br were analyzed (Figure 2C). Importantly, NTA did not detect any signal even when solutions containing 10 mg/mL Br (i.e. 1.0 g/dL), which is over 400-folds the amount of bilirubin measured in Gunn rat sera [4.1 ± 1.25 mg/dL (n = 4); CTRL rats: 0.13 ± 0.009 mg/dL (n = 9; Figure 2C)].

As consequence of red blood cells rapture, hemolysis may occur during blood collection and cytosolic content, including Hb, may be released in the serum. To evaluate whether hemolysis might contribute to NTA signal, hemoglobin was dissolved to simulate different levels of hemolysis from 0.5% to 50% (Figure 2D). Here we show that Hb becomes only detectable by NTA at concentration of 10 mg/mL (equivalent to ∼5% hemolysis) or higher. These data suggest that Hb is unlikely to contribute to NTA measurements in those samples where hemolysis is not detectable by visual inspection. Based on these findings, we concluded that protein aggregate or Br is unlikely to contribute to the quantification of EVs by NTA in non-hemolyzed crude sera. Nevertheless, it is recommended for hemolyzed specimens to be discharged.

### Separation of EVs from serum-protein by using an optimized PEG approach

PEG has been used to purify viral particles for more than 60 years [22]. It was previously shown that PEG can be also used to enrich EVs from biological fluids [23], however when we employed the method as described by Rider et al, a significant amount of serum-protein contaminant (e.g., albumin) was detected in the EVs-enriched pellets (Supplementary Figure 2). In order to reduce the quantity of contaminating proteins, a PEG titration curve was carried out. Here we show that a 6:1 (v/v) dilution of serum (or plasma) with PEG Precipitation Solution (PPS) stock resulted in the removal of >99% of soluble proteins from the EVs-enriched pellets. Notably, PPS performance is comparable to commercial EVs isolation reagents such as TEI (Figure 3A and Supplementary Figure 2). Importantly, when the enriched EVs were subjected to a second cycle of PPS isolation, no albumin was detected in the resulting pellets (Data not shown). Purity of the EVs preparation was evaluated by Western blot analysis of the exosomal marker TSG101 (Figure 3C), CD63 and HSP70 (Supplementary Figure 7) [6], which were exclusively detectable in the EVs enriched pellets, whereas the serum protein albumin was predominantly detected in the EVs-depleted supernatant (Figure 3D, Supplementary Figure 7). Overall, these data support the conclusion that the presented approach efficiently separate EVs from soluble serum proteins.

**Figure 3.**
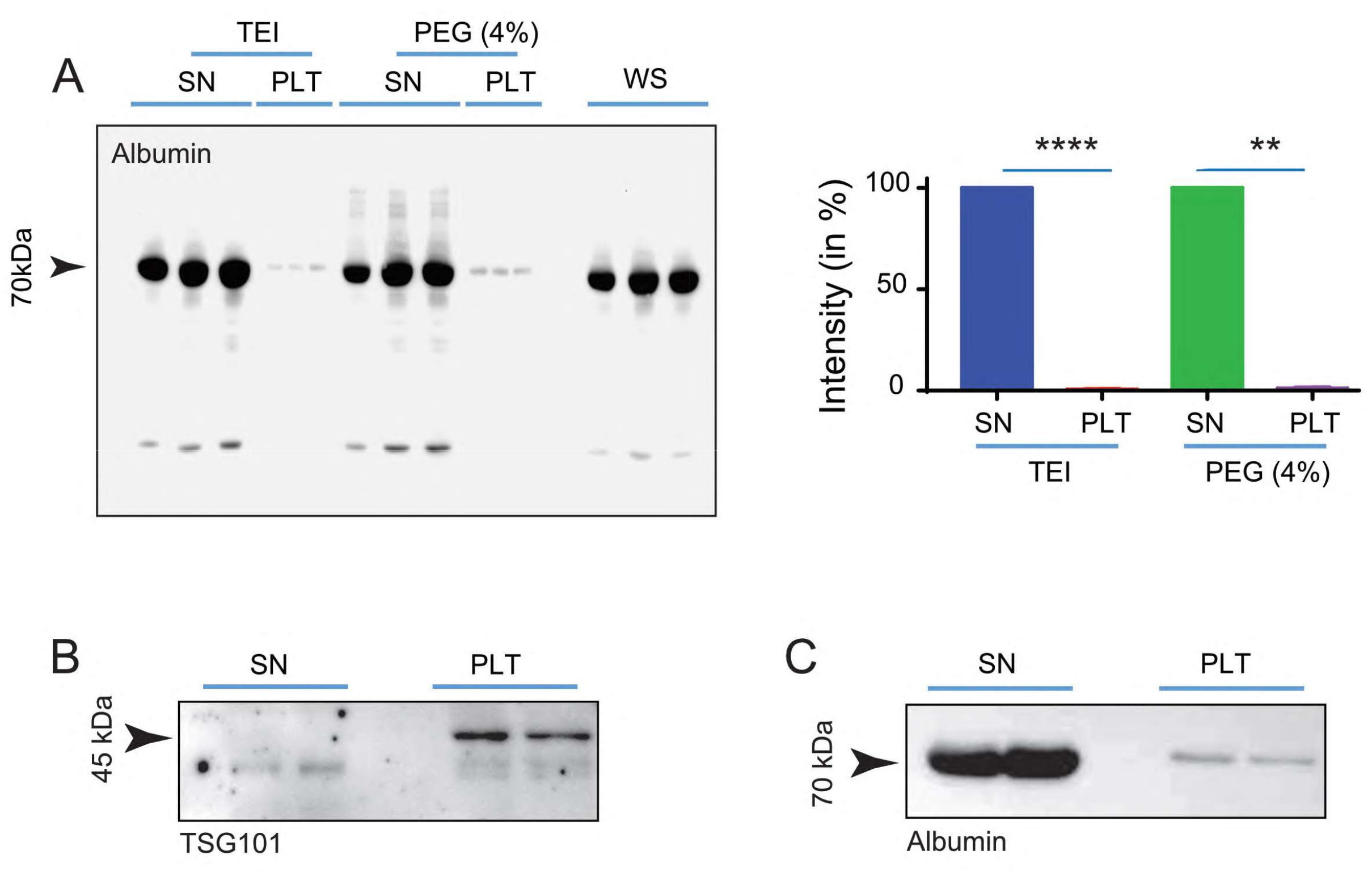
(**A**) Quantification of serum protein carry-over. TEI precipitation resulted in 0.61 % (± 0.161, n = 3), whereas PEG (4%) resulted in 1.04% (± 0.466, n = 3) carry-over of serum albumin in the pellets (PEL) (n=3). (Top panel) after 1 cycle of precipitation about 1% albumin was still detectable in the EVs-enriched pellet. (**B**) The exosomal markers TSG101 was exclusively detected in the EVs enriched pellets (PLT), on the other hand, (**C**) albumin was mainly detected in the EVs-enriched pellet (SN). Statistical analyses were carried out in GraphPad using two tailed unpaired t-tests for comparison of two groups. p ≤ 0.01, **; p ≤ 0.0001, ****.

### Evaluation of extracellular vesicles integrity and functionality

To assess the integrity of PPS-isolated EVs, measurement of cellular uptake of fluorescently labeled EVs was performed. For this purpose, EVs were fluorescently labeled with either Syto RNASelect (FITC-label) or Cell-Mask marker (PE-label). Effective EVs labeling was assessed with a BD FACSAria III flow cytometer (FC). Because the FC in use in our laboratory lacks dedicated software to quantify the size and number of nano-sized particles, the following empirical approach was used (previously described in [24]). For this purpose, fluorescently labeled particles of defined size (100 nm, 200 nm and 600 nm), were loaded on the FC, enabling the establishment of a 200 nm gate, generated with the 200 nm FITC-labeled beads (Figure 4A and 4B). Next, sera containing labeled EVs were loaded on the FC and the 200 nm gate was used to visualize both Syto RNASelect (Figure 4C) and Cell-Mask (Supplementary Figure 6) labeled EVs.

**Figure 4.**
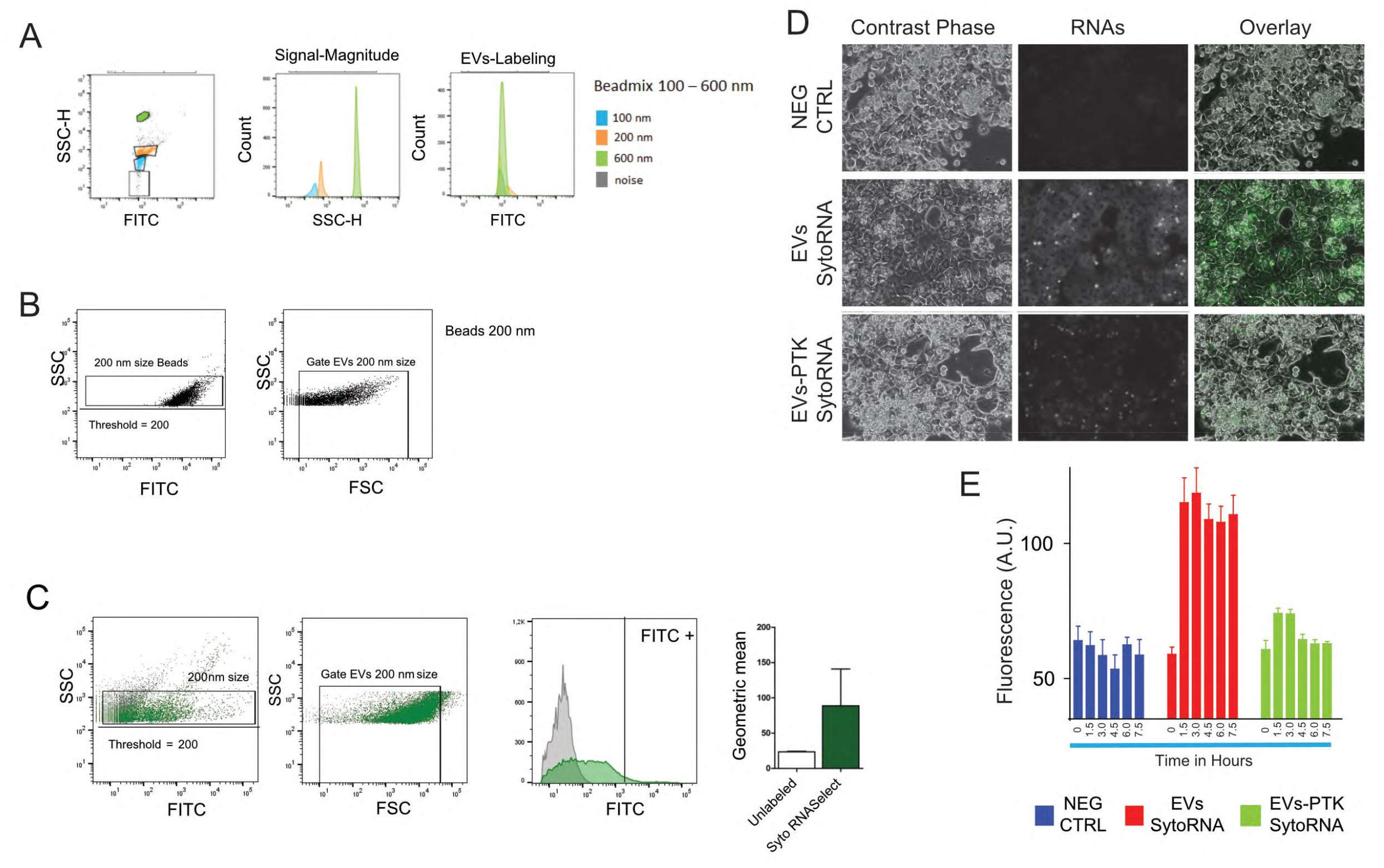
(**A**) Fluorescently labeled particles of defined size (100 nm, 200 nm and 600 nm), were loaded on the BD FACSAria III flow cytometer (FC), and (**B**) EVs of 200 nm in size were used to establish of a 200 nm gate. (**C**) Sera containing Syto RNASelect (FITC-label) labeled EVs were loaded on the FC and the 200 nm gate was used to visualize and quantify the number and fluorescence of Syto RNASelect labeled EVs. (**D**) Uptake of Syto RNASelect labeled RNA associated to EVs as acquired by fluorescence microscopy after 24 hours of incubation. (**E**) Quantification of relative fluorescence measured from primary rat hepatocytes by uptake of Syto RNASelect-labeled with or without PTK digestion EVs. Fluorescence was measured before EVs addition (time 0) and every 90 minutes for five times. Data represent average fluorescence ± SD (n = 8).

Next, sera containing fluorescently labelled RNAs were either digested with PTK or mock digested before PEG-mediated precipitation. EVs-enriched pellets were then dissolved in serum-free culture medium and supplemented to rat PCs in culture. The uptake of EVs carrying fluorescent RNAs was visualized by microscopy (Figure 4D) and fluorometric analysis was used to quantify uptake of fluorescently labeled-RNAs (Figure 4E). The observed increase in fluorescence measured in PCs incubated with intact EVs indicates that fluorescent RNAs were transferred from the EVs to the cytosol of these cells, whereas cells incubated with PTK-digested (or negative control samples) did not show any significant increase of the fluorescence in PCs. These data support the conclusion that PEG isolation does not alter EVs integrity, and these EVs are still suitable material for downstream processing/analysis.

### Analysis of miRNA expression in EVs isolated from patients with liver diseases and aged animals

In the last decade, cell free miRNAs in body fluids have emerged as a new class of biomarkers for various diseases including liver diseases [25, 26]. Cell free miRNAs are both found associated to EVs and RNA binding proteins (RBPs). Notably, electropherogram of RNAs isolated from EV-enriched pellets and EV-depleted supernatants display distinctly different RNA profiles (Figure 5A), which underscores the importance of separating the two sets of RNAs as they may possess different physiological functions.

**Figure 5.**
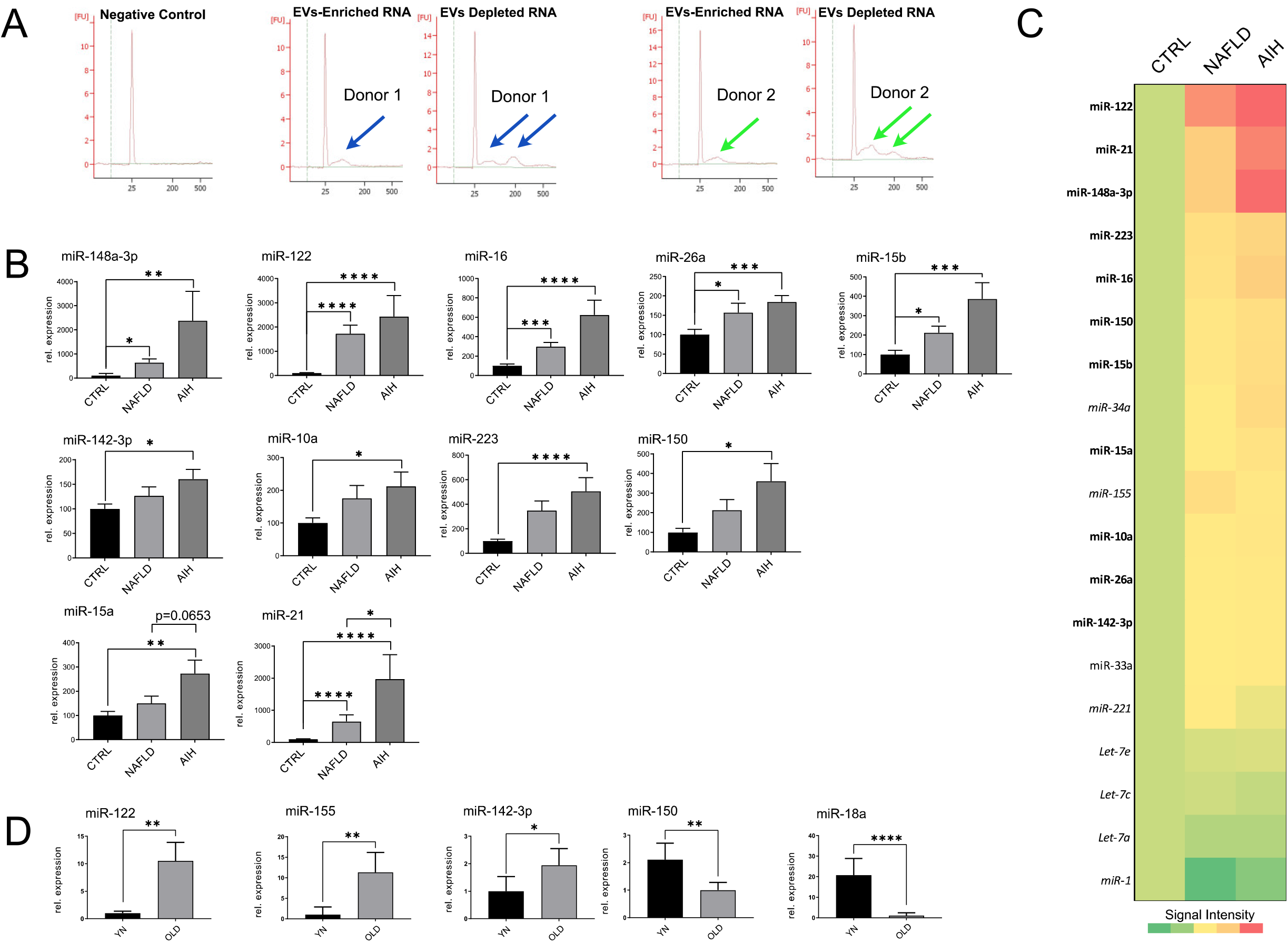
(**A**) Electropherograms of RNAs isolated from EV-enriched pellets and EV-depleted supernatants display distinctly different RNA profiles. (**B**) miQPCR was used to quantify the levels of miRNAs isolated from EVs-enriched pellets from liver-diseased patients. EVs were isolated from the sera of NAFLD patients (n = 11), AIH patients (n = 7) and healthy donors (CTRL; n = 11). (B upper panel) The expression of seven miRNAs was found significantly increased in both AIH and NAFLD vs. CTRL. (**B middle panel**) miR-142-3p, -10a and -223 expression was found to be significantly elevated in AIH vs. CTRL. (**B lower panel**) miR-150, -15a and -21 expression was found to be significantly elevated in AIH vs. NAFLD. (**C**) Analysis of selected miRNAs in EVs isolated from the sera of young (2-months-old) and old (22-months-old) rats. Statistical analyses were carried out in GraphPad using one-way ANOVA and two tailed unpaired t-tests for comparison of two groups. p ≤ 0.05, *; p ≤ 0.01, **; p ≤ 0.001, ***; ≤ 0.0001, ****.

To assess whether PPS-isolated EVs are suitable material for the identification of potential disease-associated biomarkers, a group of 11 miRNAs (Supplementary Figure 5) was selected through the mining of the publicly available study GSE113740 [26] that contains the profiling of cell-free miRNAs circulating in sera of patients with chronic hepatitis, liver cirrhosis and HCC (Supplementary Figure 5). Of the selected miRNAs, a total of five miRNAs (i.e., miR-15b, -16, 26a, -122 and -148a) were found to be significantly up-regulated in EVs isolated from AIH and NAFLD patients compared the CTRL groups (Figure 5B, top panel). The expression of another three miRNAs (i.e., miR-142-3p, -10a, and -223, Figure 5B, middle panel) was found significantly upregulated in EVs isolated from AIH patients compared to the healthy donors, whereas the expression of another three miRNAs (i.e., miR150, -15a, and -21, Figure 5B, lower panel) was found significantly upregulated in AIH vs NAFLD, AIH vs CTRL and NAFLD vs CTRL, respectively. Further studies in larger cohorts will be needed to effectively validate if this panel of miRNAs represent a liver-disease based biomarker signature with diagnostic or prognostic value.

We have previously shown that the livers of aging rats exhibit a decrease in individual extracellular matrix proteins as well as integrins when compared to young rats [27]. Furthermore, hepatic stellate cells from aged rats show a senescence-associated secretory phenotype and lowered expression of matrix proteins and growth factors. In this study we found that the distribution (e.g., size and number) of EVs circulating in the blood of older and younger animals are significantly different, closely mimicking the distribution of EVs as observed in patients with liver diseases (Figure 1). Now we show that the levels of several miRNAs normally found increased in patient with liver diseases (i.e., miR-122, -155, and -142-3p UP and miR-15- and -18a DOWN, Figure 5) are also significantly modulated in the EVs circulating in older rats compared to younger animals (Figure 5C). It would be interesting to extend this analysis to assess whether these differences in EVs distribution and miRNAs levels may actually reflect a signature of healthy aging or instead emphasize conditions of cellular senescence and inflammation.

### Luminex-based measurement of cytokines in EV-enriched pellets and EVs-depleted supernatants

Cytokines are important modulators of immune function and inflammatory responses that function as soluble factors mediating cell-cell communications in multicellular organisms. It has been reported that cytokine secretion occurs both classically and through encapsulation in EVs [28, 29]. One of the general aims of our work is the identification of early, non-invasive diagnostic biomarkers of cholangiocarcinoma (CCA). CCA has recently shown increasing mortality rates [30], therefore, there is an urgent need for biomarkers for early diagnosis of CCA. Interestingly, cytokines are among the emerging diagnostic and prognostic biomarkers in CCA [31] and IL-6, IL-8 and granulocyte-macrophage colony-stimulating factor (GM-CSF) have been found to be increased in CCA patients [32]. In order to benchmark the PEG-mediated isolation method with respect to the possibility of analyzing cytokines in EVs, the following pilot study was conducted. EVs were isolated from sera of 16 patients with CCA and from 5 healthy donors, and a Luminex 8plex was used to measure cytokines in whole sera (WS) and lysed EVs (LyEVs). Notably, differences were observed in both the type and range of cytokines detected (Figure 6A, 6B and Supplementary Table 3). While for certain cytokines the analysis of both WS and LyEVs material generated comparable results (i.e., IL-8 and IL-6), for other the analysis LyEVs material appear to provide more information. For instance, the levels of IL-2 were identified as significantly increased in the CCA group only in the samples prepared by the LyEVs approach. Moreover, GM-CSF and INFγ were exclusively detected in the samples prepared according to the LyEVs approach, although no significant differences were found between the CCA and the control groups. Overall, our data indicate that PEG-mediated isolation of EVs is compatible with cytokine analysis by Luminex. Furthermore, the data presented here suggest that (for the cytokines included in the 8plex) the LyEVs approach might be a better source of material for the discovery of diagnostic biomarkers in CCA.

**Figure 6.**
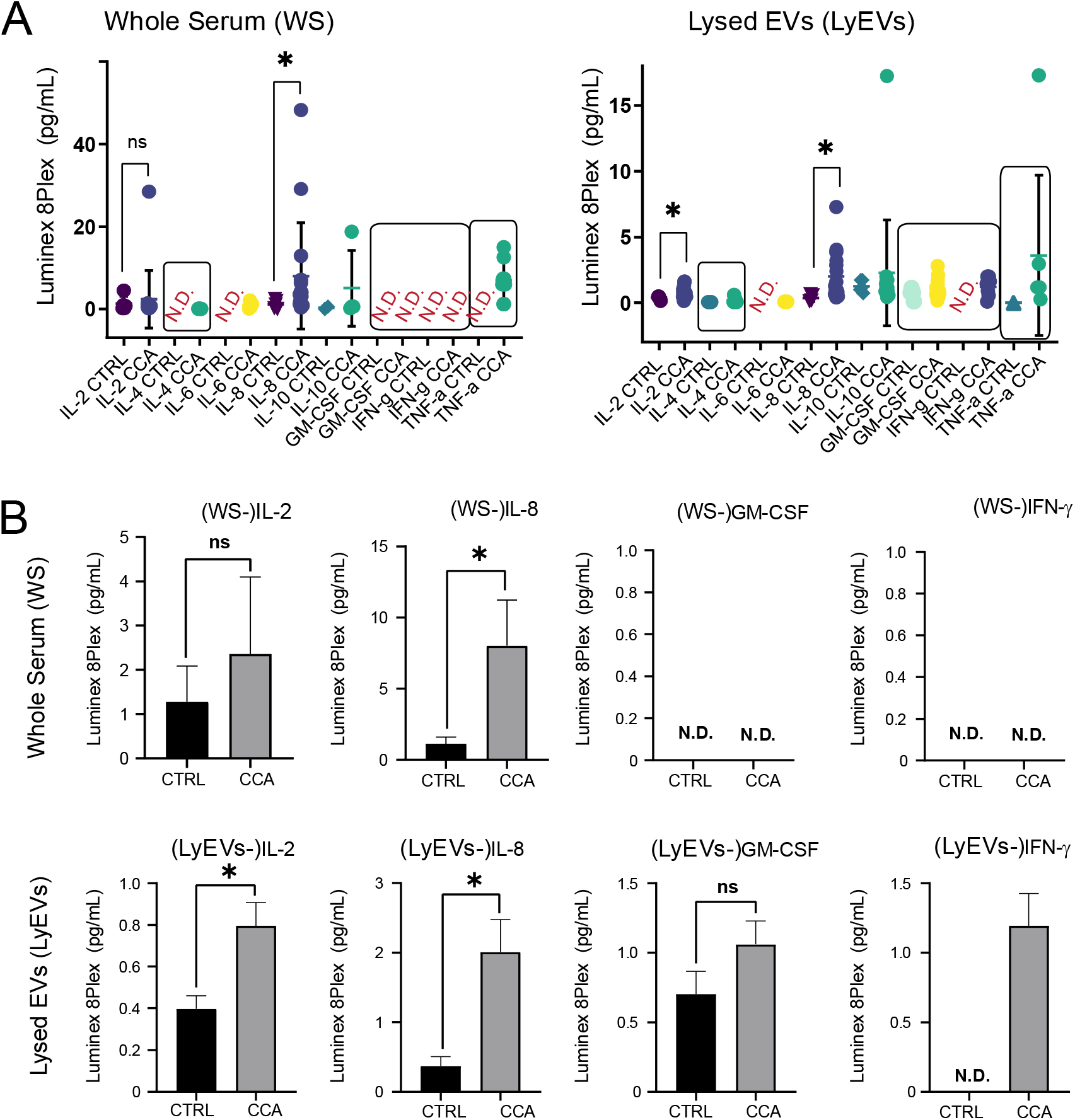
(**A**) Quantification of cytokines in sera from the CCA cohort and healthy controls (CTRL) by 8plex Luminex in whole sera (WS, left panel) and in lysed EVs (LEVs, right panel). (**B**) Side by side representation of the levels of the selected cytokines in WS (top panel) and LEVs (lower panel). Statistical analyses were carried out in GraphPad using two tailed unpaired t-tests for comparison of two groups. p ≤ 0.001, **; p ≤0.01, *.

## Discussion

Compelling evidence indicates that EVs play relevant roles in cell-cell communication and may contribute to the pathogenesis of human diseases. Numerous evidences show that isolation of EVs by existing methodologies may results in considerable carry-over of contaminants in the form of serum proteins and lipoproteins [33, 34]. Therefore, enrichment of “cleaner” EVs requires the combination of two or more isolation methods, including several control steps. While this strategy is a must when characterizing different subpopulation of EVs, it renders the task of characterizing EVs (e.g., measuring their size and number) a daunting task. In order to assess if EVs can be directly measured in crude serum by NTA an extensive number of control experiments was performed, and based on these experiments we conclude that serum-proteins, Br or Hb do not contribute to NTA measurements. These conclusions are corroborated by the observation that NTA detects no signals in EVs-depleted sera, which contains ≥99% of serum proteins. We show that NTA analysis of EVs in sera of patients with liver disease identified significant differences in both EVs size and number compared to the control group (Figure 1A). Interestingly, similar changes in EVs distribution were also observed in aged compared to younger animals (Figure 1B) and an animal model for bilirubinemia (Figure 1C). These finding are in agreement with the knowledge that inflammation might influence the machinery responsible for EVs biogenesis or secretion [14]. Hirsova et al. showed that primary hepatocytes incubated with lysophosphatidylcholine (LPC, lipotoxic compound) secrete a significant higher number of EVs [35], while Kakazu et al. demonstrated that stimulation of immortalized mouse hepatocytes with palmitic acid increase EVs secretion compared to the vehicle alone [36]. We have shown that administration of inflammatory cytokines to hepatic stellate cells (HSCs) influences both the amount and size of secreted EVs [37]. To the best of our knowledge, the presented work is the first study investigating EVs in patients with chronic liver diseases, and it supports the conclusion that analysis of EVs in crude serum might provide prognostic or diagnostic information. In this regard, we have recently shown that EVs sizes represent a novel prognostic marker in patients receiving transarterial chemoembolization (TACE) for primary and secondary hepatic malignancies [38].

In this study, we also established a cost-effective method to separate EVs from serum proteins (Figure 3) and benchmarked it against the identification of known serum miRNA biomarkers for liver diseases [25, 26]. In agreement with published literature (Supplementary Figure 5, [26]), we show that the levels of several EVs-associated miRNAs were significantly increased in patients with liver diseases but not in the control group (Figure 5B). Specifically, miR-142-3p, miR-10a, miR-150 and miR-223 were significantly higher only in EVs isolated from patients with AIH compared to healthy donor. Furthermore, the levels of miR-15a and miR-21 were found significantly increased in EVs circulating in AIH compared to NALFLP patients. Future studies in large multi-center cohorts will be required to establish whether the combination of these miRNAs could be used as reliable diagnostic or prognostic markers in liver diseases. Interestingly, strikingly similarity in EVs distribution and miRNAs levels were also observed when comparing young with aged rats (Figure 1B and Figure 5C). These data emphasize the importance of matching donor age, as some of the observed differences may actually indicate difference in age rather than indicating ongoing disease. Beside miRNAs, EVs transport other types of biological active molecules. To this end, it was shown that cytokines are present in EVs [28, 29], and that some of the EVs-encapsulated cytokines are not detectable by standard assays [29]. As cytokines are emerging candidate biomarkers in human diseases including cholangiocarcinoma [30-32], we investigated whether PEG-enriched EVs could be a suitable source for analyzing EVs-encapsulated cytokines. Here we show that not only our method is compatible with cytokine detection and measurement by Luminex (Figure 6), but also that cytokine contained in EVs displayed a better dynamic range compared to cytokines circulating in whole serum. Furthermore, Luminex was able to detect INFγ and GM-CSF cytokines only in LyEVs samples.

In conclusion, we presented an optimized and easy-to-use workflow that enables qualitative and quantitative characterization of EVs in complex biological fluids and their enrichment for subsequent analysis, including biomarker and functional characterization.

## Authors Contributions

MP, carried out experiments, analyzed the data and wrote the manuscript. SL, performed Luminex analysis of CCA samples. CK, provided and analyzed the Gunn and aged rats. VK, selected the patients from the Biobank. MW, performed FC analysis of labeled EVs. UPN, generated and curated the cohort of CCA patients. DS and TL, participated to research design. MC, conceived, designed, performed experiments, and wrote the manuscript.

All authors read and approved the manuscript.

## Acknowledgments

The authors are grateful to Claudia Rupprecht and Lisa Knopp for technical assistance. The authors acknowledge the Rat Resource & Research Center (P40OD011062) for providing Gunn rats.

## Abbreviations

miRNA: microRNA
EVs: extracellular vesicles
Exo: exosomes
MVs: microvesicles
AP: apoptotic bodies
NAFLD: nonalcoholic fatty liver disease
BTC: Billiary tract cancer
HCC: Hepatocellular carcinoma
CAA: Cholangiocarcinoma
HSP70: 70-kDa heat shock protein 70
TSG101: Tumor susceptibility gene 101
CD63: Cluster of differentiation 63
CD81: Cluster of differentiation 81
HSP70: heat shock protein 70 kDa
LPC: lysophosphatidylcholine
HSCs: hepatic stellate cells
PCs: hepatocytes
AIH: autoimmune hepatitis
TACE: transarterial chemoembolization
PEG: polyethylene glycol
RCF: relative centrifugal force
PPS: PEG precipitation solution
TEI: Total Exosome Isolation
PVF: particles per visual field
NTA: nanoparticle tracking analysis
FC: flow cytometry
TEM: transmission electron microscopy
DLS: dynamic light scattering
GOT: glutamic-oxaloacetic transaminase
GTP: glutamyl transpeptidase
Br: bilirubin
Hb: hemoglobin
TCA: trichloroacetic acid
NaCl: sodium chloride
PTK: Proteinase K
TX100: Triton X-100
RBPs: RNA binding proteins
CCA: cholangiocarcinoma
WS: whole sera
LyEVs: lysed EVs
IL-2: Interleukin 2
IL-6: Interleukin 6
IL-8: Interleukin 8
INFγ: Interferon gamma
GM-CSF: Granulocyte-macrophage colony-stimulating factor.

