## Supplemental Figures and Tables for "An optimized workflow for analyzing extracellular vesicles as biomarkers in liver diseases"

**Supplementary Figure 1.** Representative NTA images for the different samples included in this study.

**Supplementary Figure 2.** Increased amount of PEG increase serum protein carry over. Pooled rat sera were aliquot in identical volumes and processed for EVs precipitation either using TEI [1:5 (v/v)], or increasing amount of 4% PEG [1:5 (v/v)], 8% PEG [3:5 (v/v)] or 12% PEG [1:1 (v/v)]. Control samples (CTRL) were treated with PBS [1:5 (v/v)]. Notably, addition of 8% and 12% PEG correspond to the optimal range as published by Rider et al. (ExtraPEG). Following centrifugation at 10,000 rpm for 10 min at 4°C, vesicle-depleted supernatants (SN) were transferred to fresh reaction tubes. Vesicle-enriched pellets (PEL) were washed with PBS and dissolved in fresh PBS. Western Blot for albumin illustrating the albumin carry-over in the EVs-enriched pellets in three replicates per condition.

**Supplementary Figure 3. NTA quantification of polystyrene beads (ps110).** Calibration polystyrene beads of 110 nm (ps110) of diameter were diluted 1:250,000 in Ampuwa water to achieve between 300 to 50 particles per visual field (PVF) and measured by NTA. (**A**) Correlation of EV concentration (particles/mL) as measured in dependence of number of particles PVF in rat sera shows a high linearity. (**B**) Measured EV concentration was adjusted to the corresponding dilution factor and original EVs concentration (particles/cm$^3$) was illustrated against PVF. (**C**) Correlation between average EVs size (nm) as measured by NTA and PVF shows an increase in size estimation with lower PVF. (**D**) Box-plot representation of EVs size distribution for a certain PVF. PVFs are indicated as average (n = 5). Statistical analyses were carried out by GraphPad Prism by using either linear regression or two tailed unpaired t-tests.

**Supplementary Figure 4.** Pooled rat sera were aliquoted in identical volumes and EVs were precipitated using either TEI [1:5 (v/v)] or increasing amount of PEG (as indicated). Control samples (CTRL) were left untreated. Following EVs-precipitation, EVs-enriched pellets were washed twice in ice-cold PBS and dissolved in equal volumes of PBS and the (A) number and (B) size of EVs was quantified by NTA. These data shows that EVs precipitation by mean of TEI or PEG has a significant effect of the quantification of EVs by using NTA.

**Supplementary Figure 5.** In order to independently validate our findings on the expression of EVs-associated miRNAs in patients with liver diseases (see Figure 5), we compared our data with the miRNA expression profiles of liver diseased patients from publicly available datasets. For this purpose, miRNA expression datasets GSE113740 was downloaded and analyzed. This dataset contains non-coding RNA profiling (microarray) analyzed from the sera from 345 patients with hepatocellular carcinoma (HCC), 46 patients with chronic hepatitis, 93 patients with liver cirrhosis, and 1033 non-cancer individuals. Individual expression levels of selected miRNAs are shown. Statistical analyses were carried out by GraphPad Prism by one-way ANOVA. $p \leq 0.05$, *; $p \leq 0.01$, **; $p \leq 0.001$, ***; $p \leq 0.0001$, ****.

**Supplementary Figure 6.** (**A**) EVs isolated from rat sera were labeled with Cell-Mask marker (PE-label) and loaded on the BD FACSAria III flow cytometer with using a 200 nm gate generated by using fluorescently labeled 200 nm beads. (**B**) Number of unlabeled (left) and Cell-Mask labeled (right) EVs and their quantification.

**Supplementary Figure 7.** Detection of protein carryover (**Top panel**) and the Hsp70 and Cd63 exosomal markers (**Lower panel**) in EVs enriched pellets and EVs depleted supernatant after one cycle of precipitation. The exosomal markers Hsp70 and Cd63 were exclusively detected in the EVs enriched pellets, but not in the EVs-depleted supernatants (n = 4).

**Supplementary Figure 8.** (**A-C**) Unsupervised hierarchical clustering of the expression of miRNAs in EVs isolated from the plasma of NAFLD, AIH patients and from healthy donors (CTRL).

**Supplementary Table 1.** Biochemical parameters in Gunn and control rats

**Supplementary Table 2.** Assessment of the reproducibility of NTA measurements across different days

**Supplementary Table 3.** Cytokine measurement by Luminex in the WS and LyEVs of CCA patients

**Supplementary Table 4.** List of primers used in this study

**Supplementary Table 5.** Baseline biochemical parameters in CCA patient groups

Supplementary Figure 1

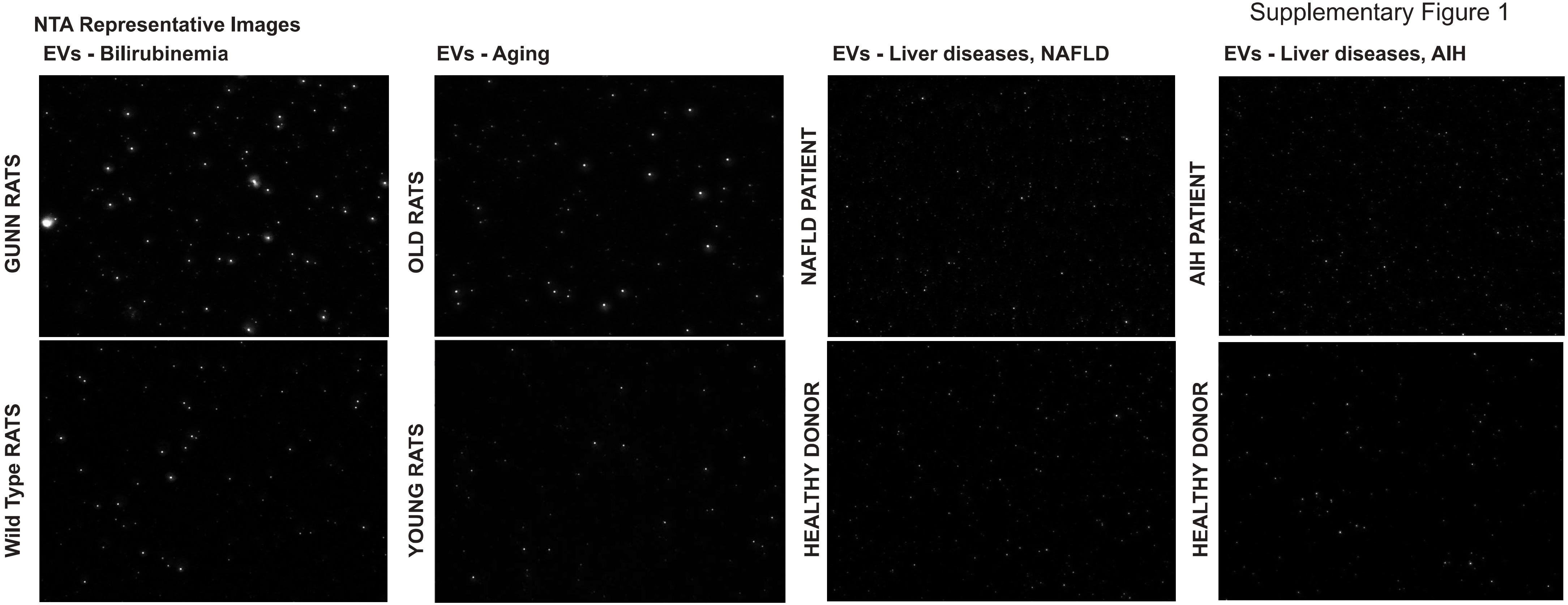

### Supplementary Figure 2

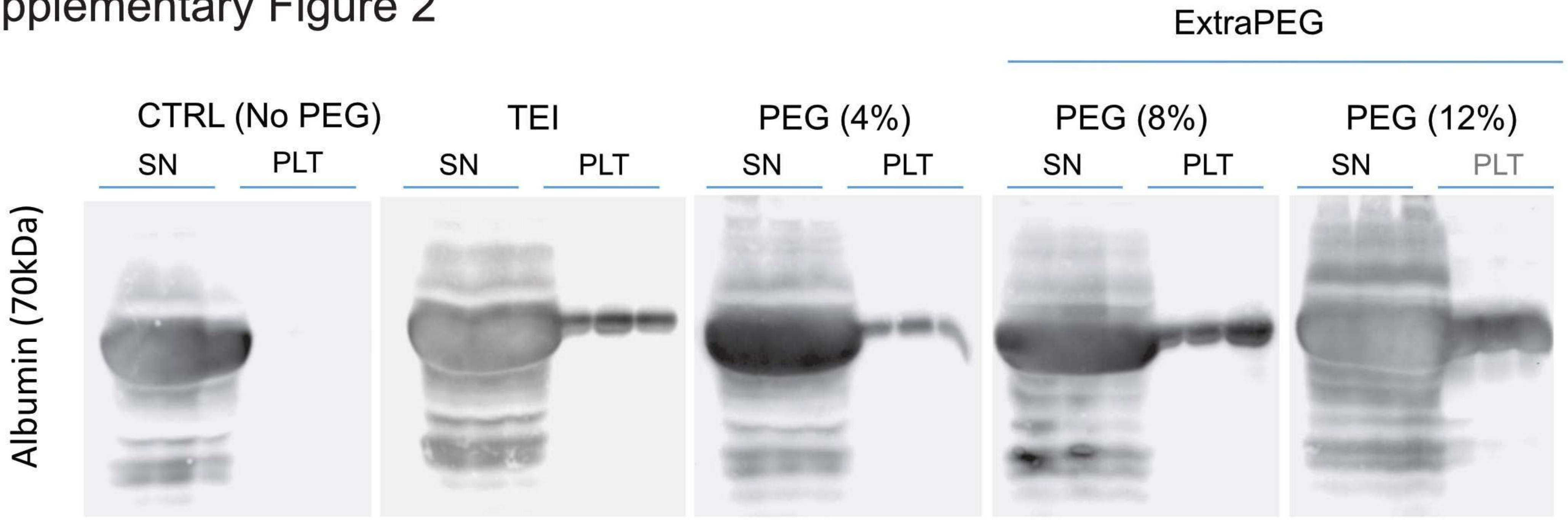

Supplementary Figure 3

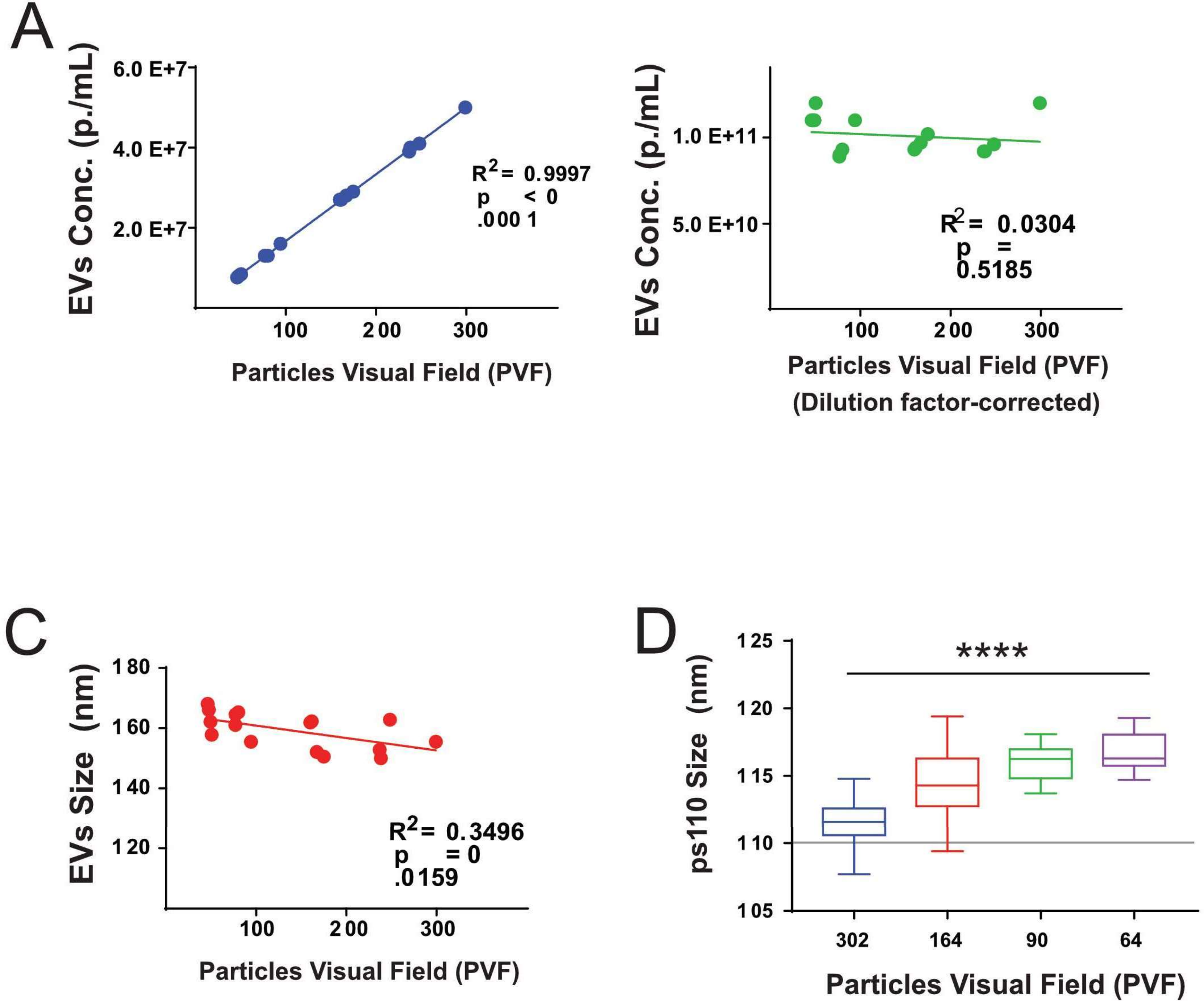

Supplementary Figure 4

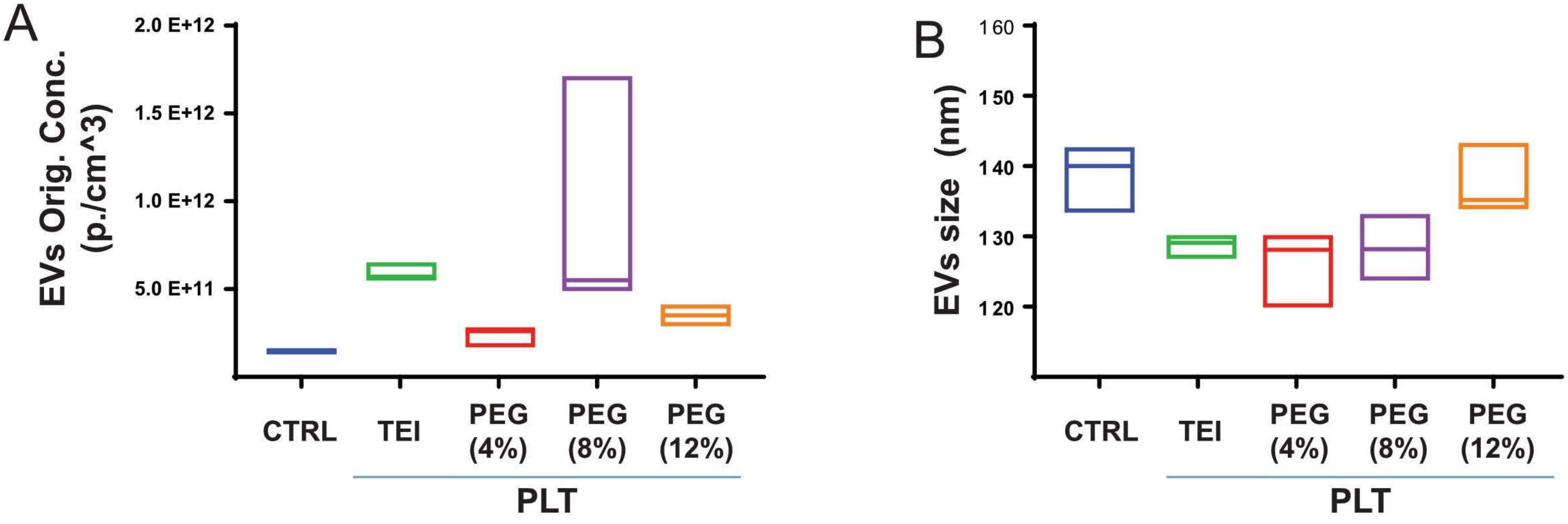

Supplementary Figure 5

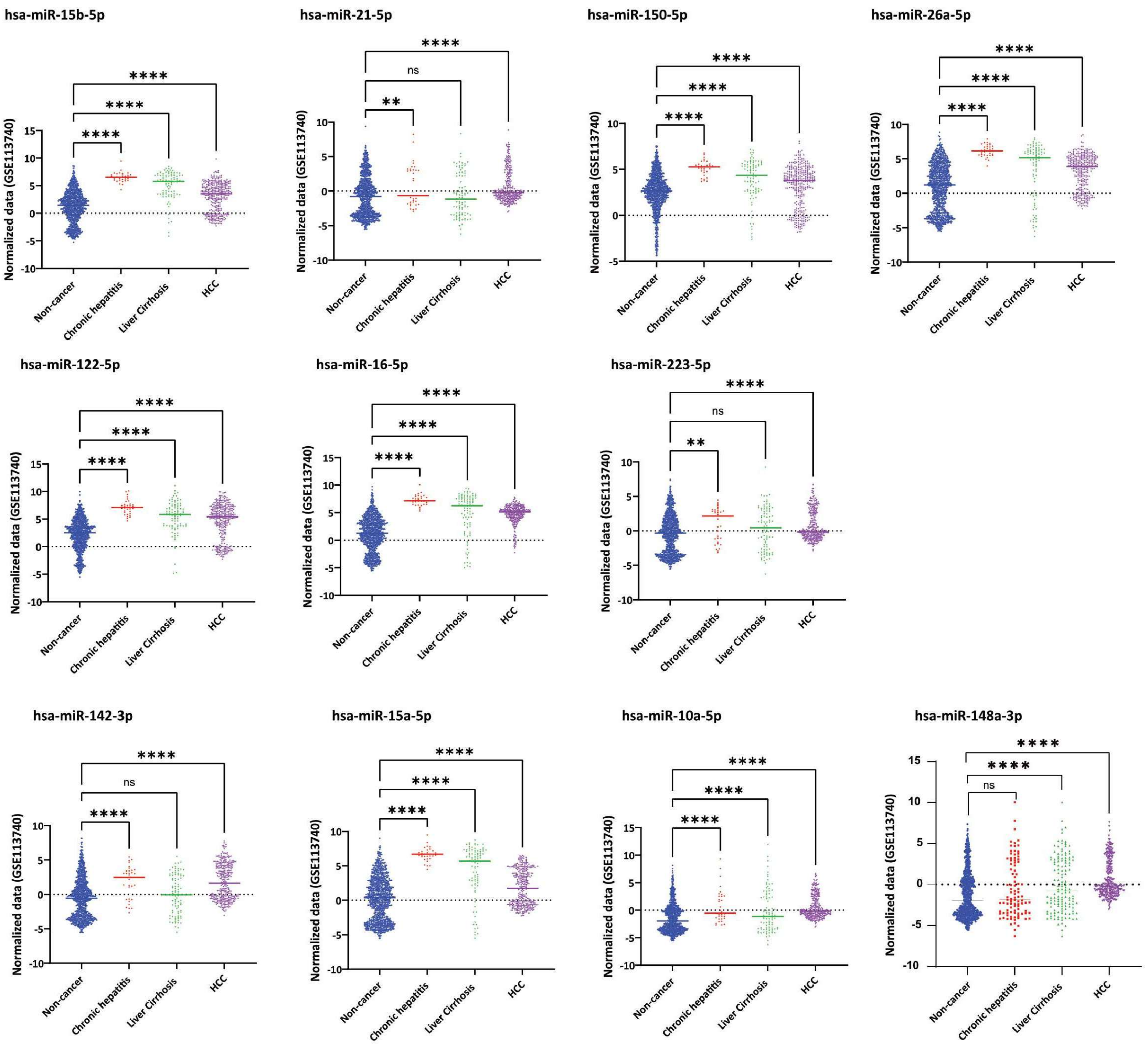

Supplementary Figure 6

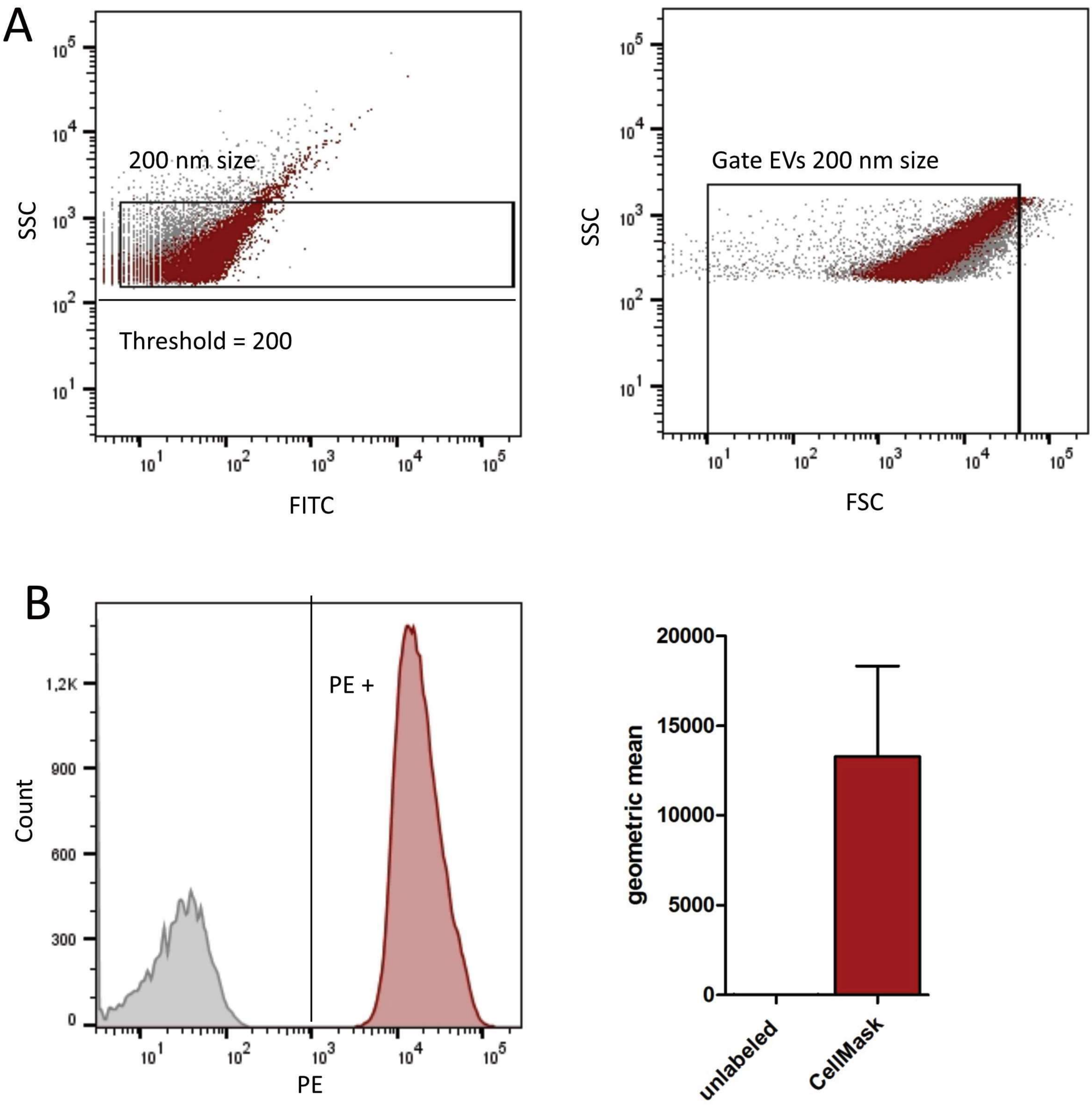

#### Supplementary Figure 7

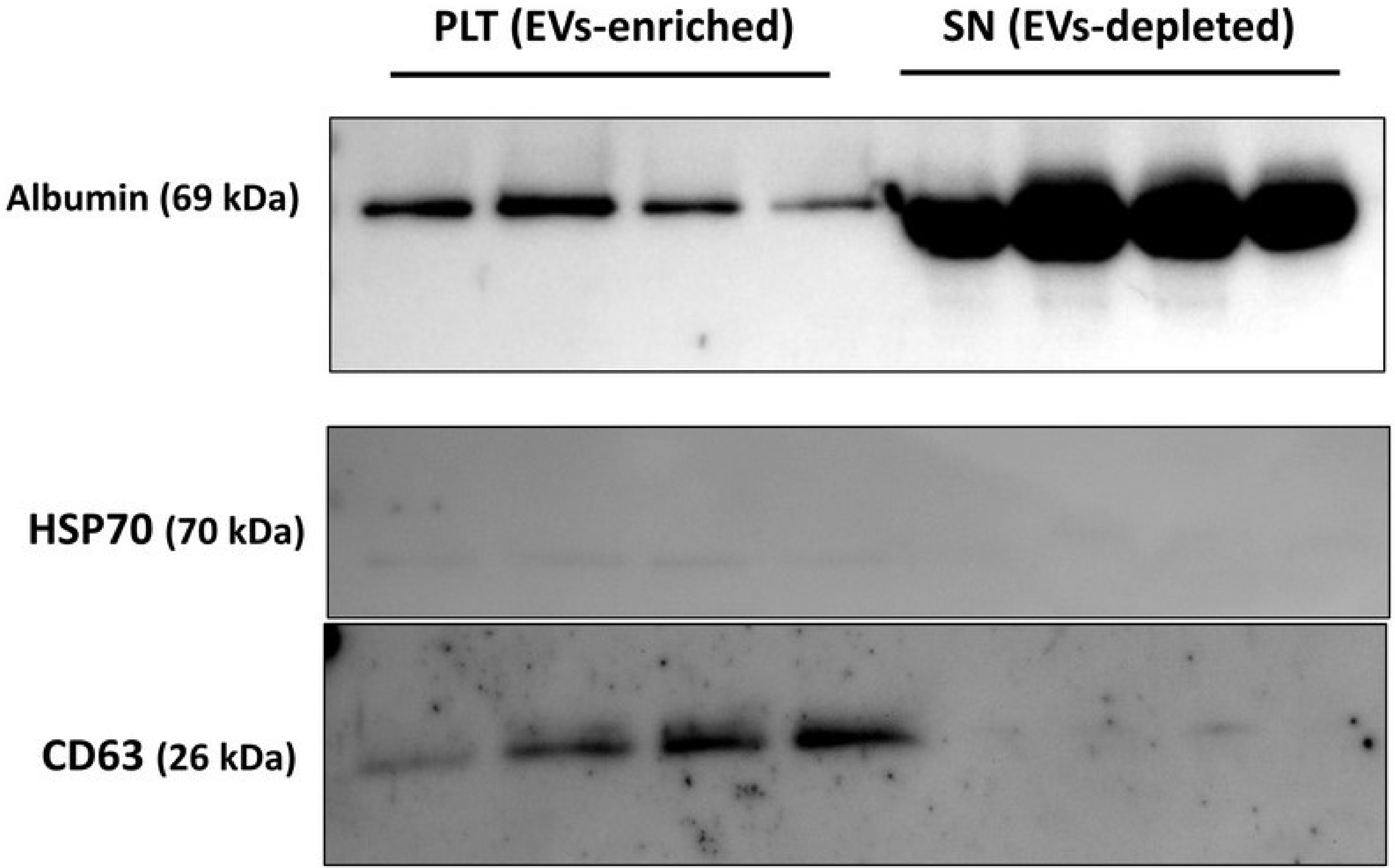

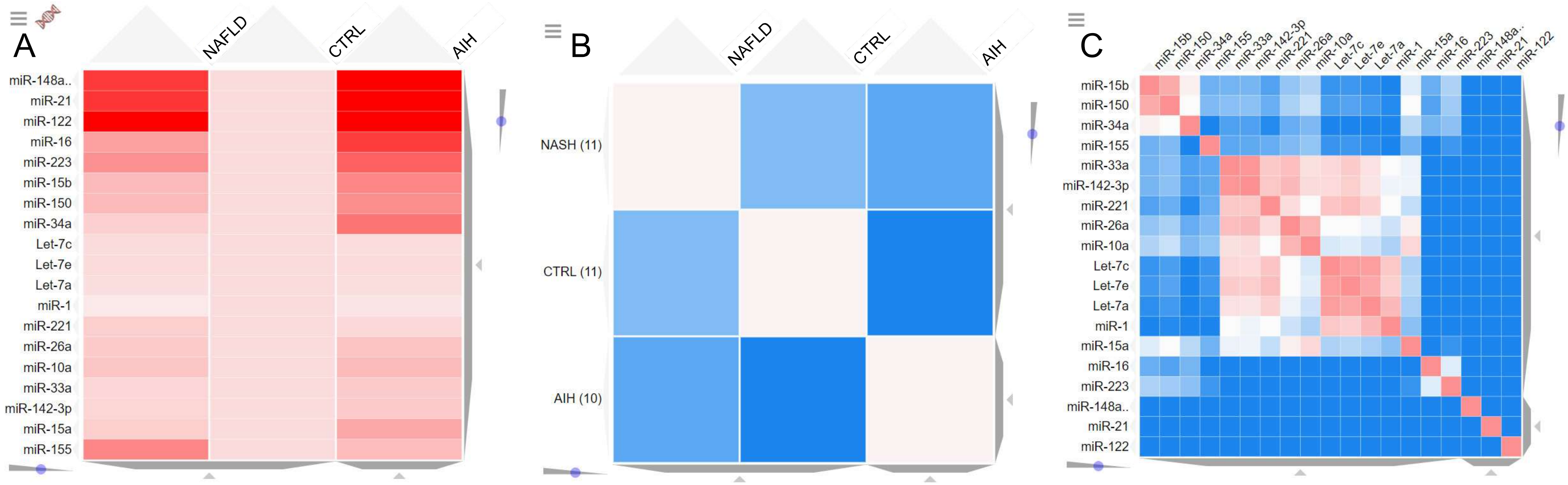


Supplementary Figure 8

**Supplementary Table1**
**Serum Gunn Rat**

| No. | Sample | T-Bil mg/dl | direct Bil mg/dl | indirect Bil mg/dl | Alb g/dl | Experiment |
|---|---|---|---|---|---|---|
| 1 | Gunn hetero 1 | 0.1 | 0 | 0.1 | 3.9 | |
| 2 | Gunn hetero 2 | 0.1 | 0 | 0.1 | 4.2 | |
| 3 | Gunn hetero 3 | 0.1 | 0 | 0.1 | 4 | |
| 4 | Gunn hetero 4 | 0.1 | 0 | 0.1 | 3.8 | |
| 5 | Gunn WT1 | 0.1 | 0 | 0.1 | 4.3 | |
| 6 | Gunn WT2 | 0.1 | 0 | 0.1 | 4.1 | |
| 7 | Gunn WT3 | 0.1 | 0 | 0.1 | 3.9 | |
| 8 | normal WT1 | 0.1 | 0 | 0.1 | 3.6 | |
| 9 | normal WT2 | 0.13 | 0 | 0.13 | 4.2 | |
| 10 | normal WT3 | 0.1 | 0 | 0.1 | 3.6 | |
| 11 | Gunn homo 1 | 5.23 | 0.32 | 4.91 | 3.5 | |
| 12 | Gunn homo 2 | 5.24 | 0.33 | 4.91 | 3.5 | |
| 13 | Gunn homo 3 | 3.72 | 0.35 | 3.37 | 4.4 | |
| 14 | Gunn homo 4 | 2.22 | 0.44 | 1.78 | 3.9 | |

| | T-Bil mg/dl | SD |
|---|---|---|
| CTRL rat | 0.11 | 0.01 |
| Gunn hetero | 0.10 | 0.00 |
| Gunn homo | 4.10 | 1.25 |
| | direct Bil mg/dl | |
| CTRL rat | 0.00 | 0.00 |
| Gunn hetero | 0.00 | 0.00 |
| Gunn homo | 0.36 | 0.05 |
| | indirect Bil mg/dl | |
| CTRL rat | 0.11 | 0.01 |
| Gunn hetero | 0.10 | 0.00 |
| Gunn homo | 3.74 | 1.30 |
| | Alb g/dl | |
| CTRL rat | 3.95 | 0.28 |
| Gunn hetero | 3.98 | 0.15 |
| Gunn homo | 3.83 | 0.37 |

**Supplementary table 2**

| Date | Name | Particle Visual Fields (PVF) | Conc. (p./mL) | Orig. Conc. (p./cm^3) | X50 (nm) | Peak ø (nm) | Span | Drift (µm/s) | Removal |
|---|---|---|---|---|---|---|---|---|---|
| 09.03.2018 | ps110.test | 279.6 | 4.60E+07 | 1.20E+13 | 111.90 | 116.30 | 0.6 | 10.6 | |
| 09.03.2018 | ps110.test | 213.4 | 3.50E+07 | 8.90E+12 | 111.00 | 113.00 | 0.7 | 9.3 | |
| 27.02.2018 | ps110.test | 224.7 | 3.70E+07 | 9.30E+12 | 108.50 | 112.00 | 0.6 | 10.6 | |
| 27.02.2018 | ps110.test | 218.9 | 3.60E+07 | 9.10E+12 | 106.60 | 110.30 | 0.7 | 7.1 | |
| 26.02.2018 | ps110.test | 227.0 | 3.80E+07 | 9.40E+12 | 109.60 | 114.10 | 0.7 | 7.6 | |
| 26.02.2018 | ps110.test | 228.2 | 3.80E+07 | 9.50E+12 | 106.70 | 110.00 | 0.7 | 7 | |
| 21.02.2018 | ps110.test | 197.9 | 3.30E+07 | 8.20E+12 | 115.20 | 117.30 | 0.7 | 6.3 | |
| 21.02.2018 | ps110.test | 234.5 | 3.90E+07 | 9.70E+12 | 112.10 | 115.30 | 0.7 | 9.8 | |
| 20.02.2018 | ps110.test | 234.2 | 3.90E+07 | 9.70E+12 | 108.70 | 112.50 | 0.7 | 8.3 | |
| 20.02.2018 | ps110.test | 200.5 | 3.30E+07 | 8.30E+12 | 108.70 | 109.30 | 0.7 | 7.2 | |
| 19.02.2018 | ps110.test | 212.3 | 3.50E+07 | 8.80E+12 | 107.00 | 108.60 | 0.7 | 6.8 | |
| 19.02.2018 | ps110.test | 183.1 | 3.00E+07 | 7.60E+12 | 116.70 | 120.90 | 0.8 | 9.7 | |
| 16.02.2018 | ps110.test | 248.9 | 4.10E+07 | 1.00E+13 | 109.40 | 111.30 | 0.7 | 10.2 | |
| 16.02.2018 | ps110.test | 261.7 | 4.30E+07 | 1.10E+13 | 110.40 | 118.10 | 0.7 | 8.8 | |
| 06.02.2018 | ps110.test | 247.7 | 4.10E+07 | 1.00E+13 | 107.70 | 114.40 | 0.7 | 6.7 | |
| 06.02.2018 | ps110.test | 242.0 | 4.00E+07 | 1.00E+13 | 108.00 | 111.80 | 0.7 | 4.8 | |
| 02.02.2018 | ps110.test | 226.1 | 3.80E+07 | 9.40E+12 | 113.70 | 118.10 | 0.7 | 9 | |
| 02.02.2018 | ps110.test | 219.5 | 3.60E+07 | 9.10E+12 | 112.70 | 116.00 | 0.7 | 8 | |
| 29.01.2018 | ps110.test | 252.6 | 4.20E+07 | 1.00E+13 | 111.00 | 115.70 | 0.7 | 10.2 | |
| 29.01.2018 | ps110.test | 240.4 | 4.00E+07 | 1.00E+13 | 108.50 | 114.10 | 0.6 | 8.1 | |
| 08.05.2018 | ps110.test | 329.4 | 5.50E+07 | 1.38E+13 | 110.1 | 111.9 | 0.7 | 7.5 | |
| 08.05.2018 | ps110.test | 295.6 | 4.90E+07 | 1.20E+13 | 109.4 | 115.6 | 0.7 | 7.1 | |
| | | **Particle Visual Fields (PVF)** | **Conc. (p./mL)** | **Orig. Conc. (p./cm^3)** | **X50 (nm)** | **Peak ø (nm)** | | | |
| | **Average** | **237.19** | **3.93E+07** | **9.81E+12** | **110.16** | **113.94** | | | |
| | **StDev** | **33.07** | **5.56E+06** | **1.38E+12** | **2.69** | **3.19** | | | |

**Note,** Particle Matrix indicates the PVF as "Av. No of Particles"

**Supplementary Table3**

| Type | Name | Well | Hu IL-2 (38) | Hu IL-4 (52) | Hu IL-6 (19) | Hu IL-8 (54) | Hu IL-10 (56) | Hu GM-CSF (34) | Hu IFN-g (21) | Hu TNF-a (36) | |
|---|---|---|---|---|---|---|---|---|---|---|---|
| X1 | WS.01 | B2 | 0.15 | | | 1.12 | 0.01 | | | | Whole Serum |
| X2 | WS.02 | C2 | 0.75 | | | 0.02 | | | | | Measured 50 uL |
| X3 | WS.03 | D2 | 0.15 | 0.05 | | 0.17 | | | | | undiluted |
| X4 | WS.04 | E2 | 4.45 | | | 1.92 | 0.57 | | | | |
| X5 | WS.05 | F2 | 0.90 | | | 2.40 | | | | 5.99 | |
| | **AVE** | | **1.28** | **0.05** | **#DIV/0!** | **1.13** | **0.29** | **#DIV/0!** | **#DIV/0!** | **5.99** | |
| | **SD** | | **1.80** | **#DIV/0!** | **#DIV/0!** | **1.05** | **0.40** | **#DIV/0!** | **#DIV/0!** | **#DIV/0!** | |
| X6 | WS.06 | G2 | 0.61 | | 1.17 | 12.90 | 0.29 | | | 1.17 | <--- This side |
| X7 | WS.07 | H2 | 0.46 | 0.12 | | 1.24 | | | | | |
| X8 | WS.08 | A3 | 0.15 | | | 0.83 | | | | | |
| X9 | WS.09 | B3 | 0.75 | | | 6.30 | | | | 1.17 | |
| X10 | WS.10 | C3 | 0.75 | 0.05 | 1.99 | 7.12 | | | | 7.38 | |
| X11 | WS.11 | D3 | 0.15 | | | 3.05 | | | | 7.38 | |
| X12 | WS.12 | E3 | 0.61 | | 0.13 | 3.05 | 0.57 | | | | |
| X13 | WS.13 | F3 | 1.32 | 0.05 | | 1.04 | | | | 14.95 | |
| X14 | WS.14 | G3 | 0.46 | 0.05 | | 1.28 | | | | | |
| X15 | WS.15 | H3 | 0.46 | | | 0.41 | | | | | |
| X16 | WS.16 | A4 | 0.75 | | | 48.24 | | | | 5.99 | |
| X17 | WS.17 | B4 | 1.04 | | | 29.12 | 0.15 | | | 5.99 | |
| X18 | WS.18 | C4 | 0.46 | | | 4.18 | | | | | |
| X19 | WS.19 | D4 | 0.31 | | 0.53 | 4.02 | | | | | |
| X20 | WS.20 | E4 | 28.46 | | | 3.53 | 18.75 | | | 5.99 | |
| X21 | WS.21 | F4 | 1.04 | | | 1.91 | 0.57 | | | 12.54 | |

| Type | Name | Well | Hu IL-2 (38) | Hu IL-4 (52) | Hu IL-6 (19) | Hu IL-8 (54) | Hu IL-10 (56) | Hu GM-CSF (34) | Hu IFN-g (21) | Hu TNF-a (36) | |
|---|---|---|---|---|---|---|---|---|---|---|---|
| X43 | EVs.01 | D7 | 0.46 | | | 0.57 | 0.72 | 0.76 | | | Evs-enriched |
| X44 | EVs.02 | E7 | 0.46 | 0.05 | | 0.64 | 1.15 | 0.76 | 0.08 | | Diluted in 50uL PBS |
| X45 | EVs.03 | F7 | 0.46 | | | | 1.73 | 0.11 | | | +48 uL LysBuf +2uL BSA |
| X46 | EVs.04 | G7 | 0.15 | 0.05 | | 0.17 | 1.73 | 1.12 | | 2.95 | Measured 50 uL |
| X47 | EVs.05 | H7 | 0.46 | | | 0.10 | 1.00 | 0.76 | | | |
| | **AVE** | | **0.40** | **0.05** | **#DIV/0!** | **0.37** | **1.27** | **0.70** | **0.08** | **2.95** | |
| | **SD** | | **0.14** | **0.00** | **#DIV/0!** | **0.27** | **0.45** | **0.37** | **#DIV/0!** | **#DIV/0!** | |
| X48 | EVs.06 | A8 | 0.75 | | | 1.28 | 1.15 | 1.22 | | | <--- This side |
| X49 | EVs.07 | B8 | 0.61 | 0.05 | | 2.88 | 1.44 | 2.10 | 0.90 | 1.17 | |
| X50 | EVs.08 | C8 | 0.15 | | | 0.88 | 0.43 | 0.76 | | | |
| X51 | EVs.09 | D8 | 0.75 | 0.09 | | 1.60 | 0.57 | 1.17 | 1.99 | 2.95 | |
| X52 | EVs.10 | E8 | 0.46 | | | 0.49 | 0.86 | 0.66 | | | |
| X53 | EVs.11 | F8 | 0.46 | | | 0.41 | 1.15 | 1.37 | | | |
| X54 | EVs.12 | G8 | 0.75 | | | 0.96 | 2.02 | 0.87 | 1.63 | 1.17 | |
| X55 | EVs.13 | H8 | 1.04 | | 0.07 | 7.28 | 1.73 | 0.43 | 1.45 | 1.17 | |
| X56 | EVs.14 | A9 | 0.46 | 0.05 | | 0.49 | 1.73 | 0.55 | 0.54 | | |
| X57 | EVs.15 | B9 | 0.75 | 0.05 | 0.07 | 2.40 | 1.15 | 1.70 | 1.99 | | |
| X58 | EVs.16 | C9 | 0.46 | | | 3.85 | 0.86 | 0.71 | | | |
| X59 | EVs.17 | D9 | 1.32 | | | 0.96 | 1.73 | 2.80 | | | |
| X60 | EVs.18 | E9 | 0.75 | 0.05 | | 0.64 | 0.86 | 0.18 | | 1.17 | |
| X61 | EVs.19 | F9 | | | | 3.21 | 1.73 | 1.07 | 0.08 | | |
| X62 | EVs.20 | G9 | 1.61 | 0.60 | | 0.80 | 17.24 | 0.60 | 0.54 | 17.29 | |
| X63 | EVs.21 | H9 | 1.61 | 0.32 | 0.07 | 4.02 | 1.73 | 0.76 | 1.63 | 0.26 | |

**Supplementary Table 4**

| Name | Sequence (5’ --> 3’) |
|---|---|
| **UPm2A** | CCCAGTTATGGCCGTTTA |
| **rno-Let-7a** | GAGGTAGTAGGTTGTATAGTTGG |
| **rno-Let-7c** | TAGTAGGTTGTATGGTTG |
| **rno-Let-7e** | TAGGAGGTTGTATAGTTG |
| **rno-miR-1** | TGGAATGTAAAGAAGTATGTATGG |
| **rno-miR-10a** | CTGTAGATCCGAATTTGTGG |
| **rno-miR-122** | GAGTGTGACAATGGTGTTTGG |
| **rno-miR-142-3p** | TGTAGTGTTTCCTACTTTATGGAG |
| **rno-miR-148a-3p** | TCAGTGCACTACAGAACTTTGG |
| **rno-miR-150** | CCAACCCTTGTACCAGTGG |
| **rno-miR-155** | TTAATGCTAATTGTGATAGGGGTG |
| **rno-miR-15a** | TAGCAGCACATAATGGTTTGTGG |
| **rno-miR-15b** | TAGCAGCACATCATGGTTTACAG |
| **rno-miR-16** | CAGCACGTAAATATTGGCGG |
| **rno-miR-18a** | GGTGCATCTAGTGCAGATAGG |
| **rno-miR-192** | TGACCTATGAATTGACAGCCG |
| **rno-miR-194** | GTAACAGCAACTCCATGTGGAG |
| **rno-miR-21** | TAGCTTATCAGACTGATGTTGAG |
| **rno-miR-29b** | TAGCACCATTTGAAATCAGTGTTG |
| **rno-miR-221** | ATTGTCTGCTGGGTTTCG |
| **rno-miR-223** | TCAGTTTGTCAAATACCCCAG |
| **rno-miR-26a** | AAGTAATCCAGGATAGGCTG |
| **rno-miR-33a** | TGCATTGTAGTTGCATTGCAG |
| **rno-miR-34a** | CAGTGTCTTAGCTGGTTGTG |
| **rno-miR-92a-3p** | CTTGTCCCGGCCTGG |
| **rno-miR-98** | AGGTAGTAAGTTGTATTGTTGG |

**Supplementary Table5**

| Type of sample | Code on 8Plex | sex | BMI | age | Tumor localisation | T | N | M | R | G |
|---|---|---|---|---|---|---|---|---|---|---|
| CCA | S.06 | male | 19.32 | 65 | intrahepatic CCA | T3 | N1 | M0 | R0 | G3 |
| CCA | S.07 | male | 26.3 | 76 | intrahepatic CCA | T3 | N1 | M1 | R0 | G3 |
| CCA | S.08 | female | 32.39 | 52 | intrahepatic CCA | T3 | N1 | M1 | R0 | G3 |
| CCA | S.09 | female | 19.15 | 49 | intrahepatic CCA | T2 | N0 | M0 | R0 | G2 |
| CCA | S.10 | female | 24.8 | 68 | intrahepatic CCA | T2 | | | R0 | G3 |
| CCA | S.11 | female | 23.03 | 75 | intrahepatic CCA | T4 | | M1 | | |
| CCA | S.12 | male | 22.6 | 54 | intrahepatic CCA | T1 | N1 | M0 | R1 | G2 |
| CCA | S.13 | female | 21.87 | 60 | intrahepatic CCA | T2 | N0 | M0 | R1 | G3 |
| CCA | S.14 | male | 22.47 | 42 | intrahepatic CCA | T2 | | M0 | R0 | G2 |
| CCA | S.15 | female | 31.18 | 42 | intrahepatic CCA | T2 | N1 | M0 | R0 | G2 |
| CCA | S.16 | female | 24.61 | 84 | intrahepatic CCA | T3 | N0 | M0 | R0 | G2 |
| CCA | S.17 | male | 27.12 | 84 | intrahepatic CCA | T1 | N0 | M0 | R0 | G2 |
| CCA | S.18 | male | 27.7 | 80 | intrahepatic CCA | T1 | N0 | | R0 | G3 |
| CCA | S.19 | female | 31.25 | 74 | intrahepatic CCA | T3 | N1 | M0 | R0 | G2 |
| CCA | S.20 | female | 21.97 | 61 | intrahepatic CCA | T3 | N0 | M0 | R1 | G3 |
| CCA | S.21 | male | 31.7 | 55 | intrahepatic CCA | T4 | N1 | M1 | R1 | |
